# HiCPEP: Efficient estimation of chromatin compartment PC1 from Hi-C covariance structure

**DOI:** 10.64898/2026.05.14.725269

**Authors:** Zhi-Rong Cheng, Jia-Ming Chang

## Abstract

Principal component analysis (PCA) of the Hi-C Pearson correlation matrix is the standard approach for identifying A/B chromatin compartments. Despite its widespread use, the relationship between the first principal component (PC1) and the underlying compartment structure remains insufficiently characterized, and computing PC1 can become computationally expensive for high-resolution Hi-C data.

Here we investigate the role of the PC1 explained variance ratio in compartment analysis and show that chromosomes with strong compartment organization typically exhibit a dominant PC1 signal. Based on this observation, we propose HiCPEP, a heuristic algorithm that estimates the sign pattern and relative magnitude of PC1 directly from the Hi-C Pearson covariance matrix without performing explicit eigenvector decomposition. The method can operate from either a dense Pearson matrix for fast approximation or a sparse observed/expected (O/E) matrix to reduce memory usage. Furthermore, because many covariance columns exhibit PC1-like patterns when the compartment signal is strong, HiCPEP can be accelerated using random sampling without substantially reducing accuracy.

Across multiple Hi-C datasets, HiCPEP consistently recovered compartment patterns with high similarity to reference PC1 vectors produced by standard PCA-based methods. Benchmark experiments show that HiCPEP achieves comparable accuracy while reducing computational cost in terms of runtime or memory usage. These results suggest that HiCPEP provides a practical alternative for efficient chromatin compartment analysis from large-scale Hi-C datasets. The HiCPEP implementation is freely available at https://github.com/ZhiRongDev/HiCPEP.

## Introduction

The Hi-C technique was first proposed by Lieberman-Aiden et al. (2009), which is adapted from the chromosome conformation capture (3C) method (Dekker et al., 2002) for analyzing the folding principle and spatial organization of the whole genome. Hi-C provides a so-called “contact matrix” representing the interaction frequency between each pair of separated genomics bins (i.e., “loci”); this map can be used to identify the chromatin compartments and topologically associated domains (TADs) (Dixon et al., 2012), which are groups of loci that interact with each other more often due to a similar chromatin status (Kalluchi et al., 2023), and to detect the CTCF loops, which bring pairs of loci that are located distantly together (Rao et al., 2014).

The “resolution” of a Hi-C matrix means “The smallest locus size such that 80 percent of loci have at least 1,000 contacts” (Rao et al., 2014), which represents the finest scale a matrix could be inspected. The first contact matrix constructed by Erez Lieberman-Aiden et al. in 2009 merely divides the chromatin into 1 megabase (1 Mb) and 100 kilobase (100 Kb) per genomics bin (Lieberman-Aiden et al., 2009). Thanks to the advancement of DNA sequencing, the resolution of the matrix in 2014 had already improved to 1 Kb resolution with 4.9 billion contacts (Rao et al., 2014), and a recent publication in 2023 even resolved the resolution to 500 bp (i.e., base-pair), produced a super large matrix with about 33 billion contacts (Harris et al., 2023).

### Chromatin compartments analysis

Hi-C contact matrices exhibit characteristic “plaid” patterns reflecting the segregation of genomic loci into two major chromatin compartments, referred to as *A* and *B* compartments. Interactions between loci within the same compartment (AA or BB) are enriched, whereas interactions between compartments (AB) are depleted. In general, A compartments correspond to open, transcriptionally active chromatin, while B compartments are associated with closed, transcriptionally repressed regions (Lieberman-Aiden et al., 2009).

The classical approach for identifying A/B chromatin compartments is based on Principal Component Analysis (PCA) applied to a locus–locus correlation matrix derived from Hi-C data. Briefly, the normalized Hi-C contact matrix is first transformed into an Observed/Expected (O/E) matrix to remove the genomic distance-dependent interaction decay (Lieberman-Aiden et al., 2009; Kalluchi et al., 2023). Because many locus pairs have no observed contacts, the resulting O/E matrix is typically sparse. Next, a Pearson correlation matrix is computed between loci based on their interaction profiles in the O/E matrix, resulting in a dense matrix that highlights the plaid compartment structure. PCA is then applied to this correlation matrix, and the first principal component (PC1) is used to assign chromatin compartments: loci with positive PC1 values are typically associated with the A compartment, while negative values correspond to the B compartment. Because the sign of eigenvectors is mathematically arbitrary, the orientation of PC1 is sometimes flipped according to genomic features such as GC content to ensure that GC-rich regions correspond to the A compartment.

This PCA-based framework has become the standard approach for chromatin compartment analysis and is widely used in Hi-C analysis pipelines. In this study, we focus on the estimation of PC1 signals for the widely used binary A/B compartment analysis. However, computing the Pearson correlation matrix and performing PCA require dense matrix operations, which can be computationally expensive or unstable when the Hi-C matrix is highly sparse.

### Related works

Several Hi-C analysis tools implement PCA-based chromatin compartment identification, including POSSUMM (Harris et al., 2023), FAN-C (Kruse et al., 2020), HOMER (Heinz et al., 2010), Cooler (Abdennur and Mirny, 2020), and Juicer (Durand et al., 2016). These tools follow the classical framework of constructing a Pearson correlation matrix from the Hi-C contact map and applying PCA to obtain the first principal component (PC1), whose sign pattern defines A/B compartments.

Despite its widespread use, the PCA-based approach has two major limitations. First, the interpretation of PC1 in chromatin compartment analysis is largely empirical. While the sign of PC1 is commonly used to assign A/B compartments, the theoretical justification for this interpretation and the role of the explained variance ratio are rarely discussed (Zheng and Zheng, 2018). Second, the method requires constructing and analyzing a dense Pearson correlation matrix, which leads to substantial computational and memory costs at high resolutions. Because the space complexity of the Pearson matrix is *O*(*d*^2^), the memory requirement grows rapidly with the number of genomic bins. For example, analyzing chromosome 1 at 1 kb resolution can require hundreds of gigabytes of RAM (Kalluchi et al., 2023), making the analysis impractical for many laboratories.

To address the computational bottleneck, several studies have proposed alternative strategies. POSSUMM (Harris et al., 2023) reduces memory usage by operating directly on the sparse O/E matrix without explicitly constructing the dense Pearson matrix. Other approaches, such as CscoreTool (Zheng and Zheng, 2018), infer A/B compartments through probabilistic models rather than PCA. However, these approaches either remain computationally demanding or deviate from the widely adopted PC1-based framework.

In this study, we revisit the role of PC1 in chromatin compartment analysis and develop HiCPEP, a fast heuristic method for estimating the sign pattern of PC1 without explicitly computing the dense Pearson matrix or performing full PCA decomposition. Our approach can operate directly on sparse Hi-C matrices and can further accelerate the estimation through random sampling. This design preserves the widely used PC1- based interpretation of chromatin compartments while significantly reducing the computational requirements of compartment analysis. The algorithmic details of HiCPEP are described in the following section.

## Methods

### Overview of HiCPEP

HiCPEP is a heuristic method for estimating the sign pattern of the first principal component (PC1) used in A/B compartment analysis, without explicitly performing full PCA on the Hi-C Pearson correlation matrix. Our method is motivated by the observation that, when the Hi-C Pearson matrix exhibits a strong plaid structure, the explained variance ratio of PC1 is typically high and PC1 aligns with the dominant column patterns of the Pearson covariance matrix. Based on this observation, HiCPEP estimates PC1 by selecting a representative high-magnitude column from the covariance matrix.

We implement HiCPEP in two modes. The first approach starts from the dense Pearson correlation matrix and provides a fast approximation of PC1 without eigen-decomposition. The second starts from the sparse observed/expected (O/E) matrix and reconstructs only the required Pearson rows and covariance columns on demand, thereby reducing memory usage for high-resolution Hi-C data. In both cases, the output is an *Estimated PC1-pattern*, whose sign can be oriented using GC content so that positive values correspond to the A compartment.

### Datasets and preprocessing

We evaluated HiCPEP using Hi-C datasets from GSE18199 (Lieberman-Aiden et al., 2009; van Berkum et al., 2010) and GSE63525 (Rao et al., 2014; Sanborn et al., 2015). For GSE18199, the Pearson correlation matrices and PC1 vectors were obtained directly from the processed files `GSE18199_binned_heatmaps.zip.gz` and `GSE18199_eigenvector_files.zip.gz`. For GSE63525, Pearson correlation matrices and PC1 vectors were generated from the `combined_30.hic` files using Juicer (Durand et al., 2016) at 1 Mb, 100 Kb, and 25 Kb resolutions for the following cell lines: `CH12-LX`, `GM12878_insitu_primary+replicate`, `HMEC`, `HUVEC`, `IMR90`, `K562`, `KBM7`, and `NHEK`.

In dense-mode HiCPEP, the Pearson correlation matrix was used as input, and the corresponding PC1 was treated as ground truth for evaluation. In sparse-mode HiCPEP, the O/E matrix was used as input. All GSE63525 matrices were KR-normalized. Before all computations, NaN entries and all-zero rows or columns were removed.

### HiCPEP from a dense Pearson matrix

Let *X* ∈ ℝ^*d* × *d*^ denote the centered Pearson correlation matrix derived from Hi-C data, and let

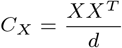

be its covariance matrix. In standard PCA, the first principal component *pc*_1_ of *C*_*X*_ is used for A/B compartment identification (details in Appendix A). When the Hi-C Pearson matrix exhibits a strong plaid structure, the explained variance ratio of *pc*_1_ is typically high, indicating that *pc*_1_ captures the dominant compartment-related variation (details in Appendix B).

Our key observation is that, under this regime, the dominant columns of *C*_*X*_ have sign patterns and relative magnitudes similar to those of *pc*_1_ (see Appendix C for details). We therefore estimate PC1 by selecting the column of *C*_*X*_ with the largest absolute column sum:

#### Algorithm 1

HiCPEP from a dense Pearson matrix

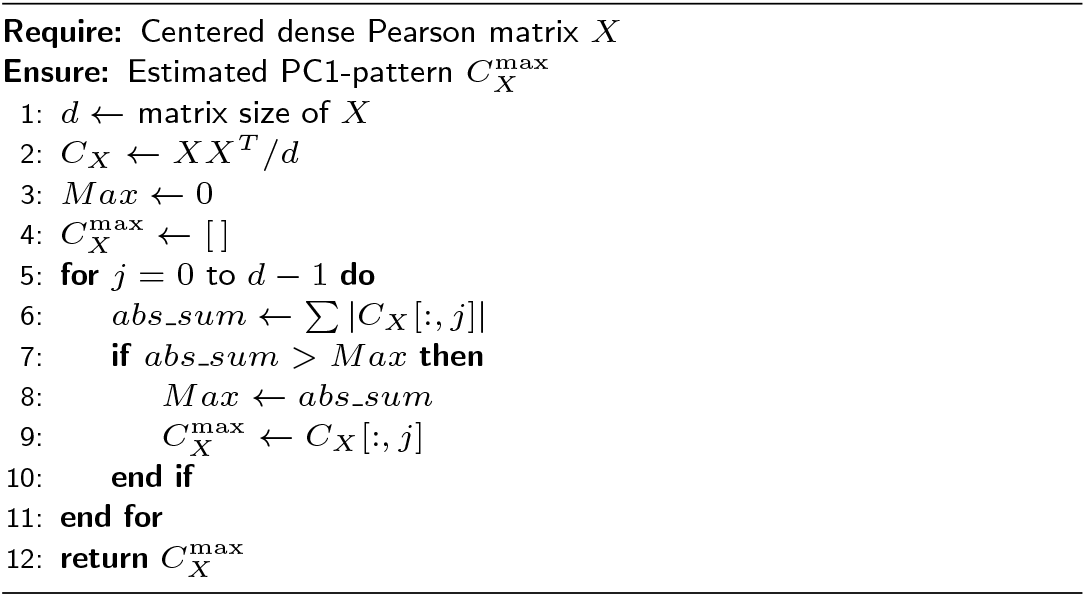

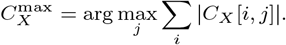

We use 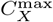 as the *Estimated PC1-pattern*. Intuitively, this column best reflects the dominant covariance structure and thus serves as a surrogate for *pc*_1_, up to a global sign flip.

The dense HiCPEP (Algorithm 1) procedure is:

1. Compute the covariance matrix *C*_*X*_ = *XX*^*T*^ */d* (Line 2).
2. For each column of *C*_*X*_, compute the sum of absolute values (Line 6).
3. Select the column with the largest absolute sum as the Estimated PC1-pattern (Lines 7 9).
4. Orient the sign using GC content so that GC-rich regions correspond to the A compartment.

The computational cost is dominated by constructing *C*_*X*_, resulting in *O*(*d*^3^) time and *O*(*d*^2^) memory.

### HiCPEP from a sparse O/E matrix

Although the dense Pearson matrix provides a convenient route to the Estimated PC1-pattern, explicitly materializing the Pearson matrix and its covariance matrix becomes impractical at high resolution due to the *O*(*d*^2^) memory requirement. To address this limitation, we derive a memory-efficient version of HiCPEP that operates directly on the sparse O/E matrix.

Let *X* denote the sparse O/E matrix, where the *i*th row is *x*_*i*_. Define the row means and standard deviations as

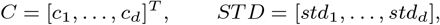

and let *I* be the all-ones row vector. The full Pearson matrix can be written as

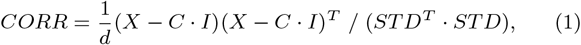

but forming this matrix explicitly would again require *O*(*d*^2^) memory.

Instead, we reconstruct one Pearson row at a time. For the *i*th row, we compute

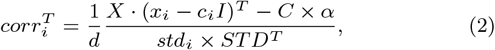

where *α* = *I* · (*x*_*i*_ − *c*_*i*_*I*)^*T*^ is a scalar. After obtaining Pearson rows on demand, we reconstruct one covariance column at a time and select the column with the largest absolute sum as the Estimated PC1- pattern. Thus, the sparse version preserves the same estimator as the dense version while avoiding construction of the full dense Pearson matrix.

The sparse HiCPEP procedure (Algorithm 2) is:

#### Algorithm 2

HiCPEP from a sparse O/E matrix

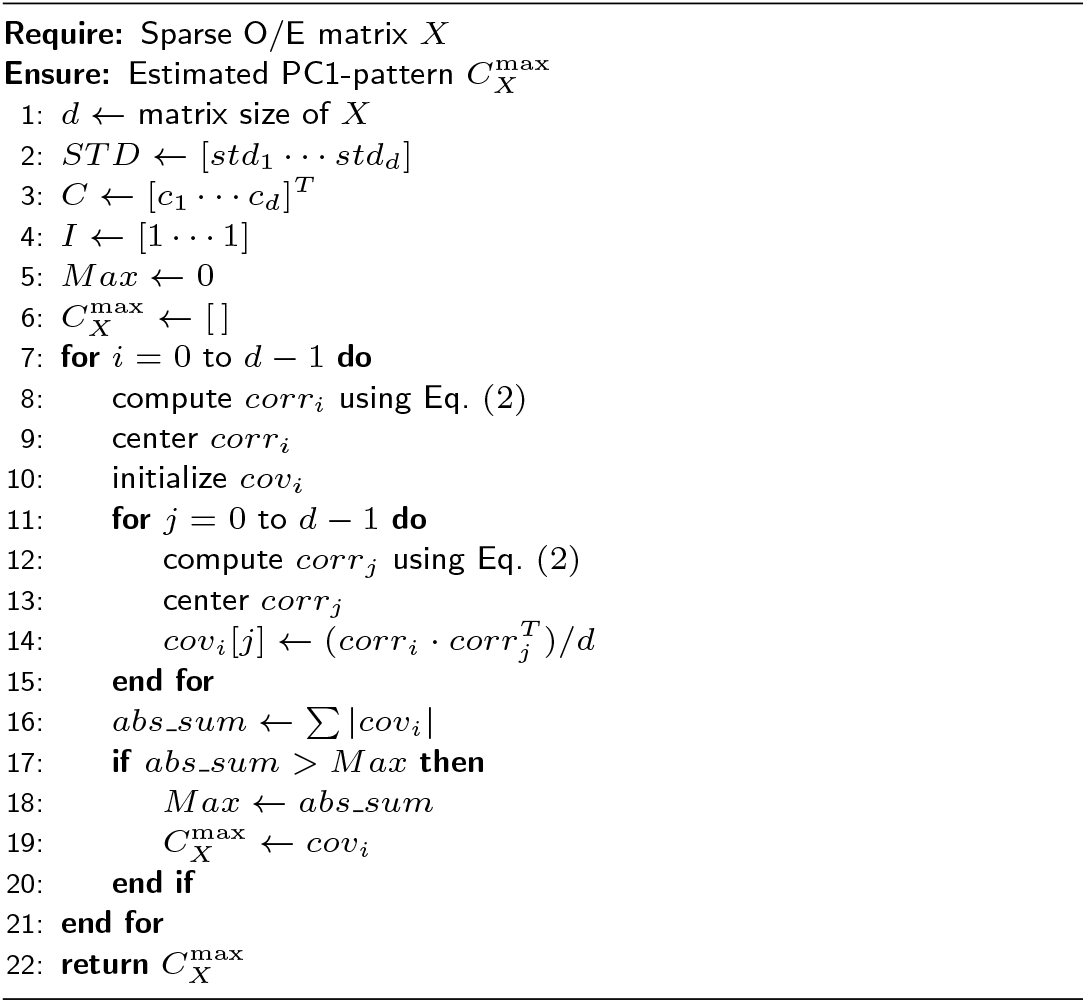

1. Reconstruct a target Pearson row *corr*_*i*_ from the sparse O/E matrix using Eq. (2).
2. Reconstruct a covariance column *cov*_*i*_ from pairwise inner products between *corr*_*i*_ and all reconstructed *corr*_*j*_ rows (Lines 11 15).
3. Compute the absolute column sum of *cov*_*i*_ (Line 16).
4. Repeat over candidate columns and select the column with the largest absolute sum as the Estimated PC1-pattern (Lines 7 21).

The time complexity is *O*(*d*^4^), whereas the space complexity is strictly less than *O*(*d*^2^) because only the sparse O/E matrix and a small number of *O*(*d*) vectors are stored. A detailed derivation is provided in Appendix D.

### Benchmark design

We benchmarked HiCPEP to evaluate both accuracy and computational efficiency. Accuracy was assessed by comparing the Estimated PC1-pattern with the ground-truth PC1 across chromosomes, cell lines, and resolutions. Efficiency was assessed in terms of runtime and peak memory usage.

#### Evaluation metric

For each chromosome, we compared the sign pattern of the Estimated PC1-pattern with that of the ground-truth PC1. Let *valid entry num* denote the number of non-NaN entries and *similar num* the number of entries with matching signs. We define

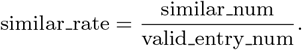

Because eigenvector signs are arbitrary, the Estimated PC1-pattern was flipped when its cosine similarity with the ground-truth PC1 was negative. For practical use, however, compartment orientation should be determined using GC content rather than the reference PC1.

#### Benchmark protocol

We evaluated dense-mode HiCPEP on Pearson matrices at 1 Mb, 100 Kb, and 25 Kb resolution, and sparse-mode HiCPEP on O/E matrices. We compared HiCPEP against Scikit-learn power iteration, POSSUMM, and Juicer.

All runtime and memory measurements were collected using `/usr/bin/time -v`, which reports the maximum resident set size (RSS). All methods were run on the same hardware platform: Intel(R) Core(TM) i7-9750H CPU @ 2.60 GHz with 16 GiB RAM.

For dense-mode HiCPEP and Scikit-learn, the benchmark command was

time −v python <script name

> For POSSUMM, the benchmark command was

time −v Rs c r i p t eig From Hic Rscript. R −v TRUE \

−n KR <hic > <output> <resolution >

> For Juicer, the benchmark command was

time −v java −jar juicer_tools_1.22.01.jar \

eigenvecto rKR <hic > “2” BP <resolution > <output>

All benchmarking scripts and configurations are available in our repository.

## Results

### HiCPEP accurately recovers PC1 compartment patterns

We first evaluated whether HiCPEP can recover the sign pattern and relative magnitude of the first principal component (PC1) from the Hi-C Pearson covariance matrix. Figure 1 shows a representative example using GSE63525 K562 chromosome 1 at 1 Mb resolution. When the column with the largest absolute column sum, 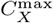, was selected as the Estimated PC1-pattern, the sign pattern closely matched the ground-truth PC1, with a *similar rate* greater than 0.98. After Z-score normalization, the estimated and ground-truth profiles also showed highly similar relative magnitudes. In contrast, selecting 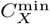 yielded a much lower *similar rate* and failed to reproduce the PC1 profile, indicating that the estimator specifically depends on the dominant covariance structure rather than on an arbitrary column.

**Figure 1.**
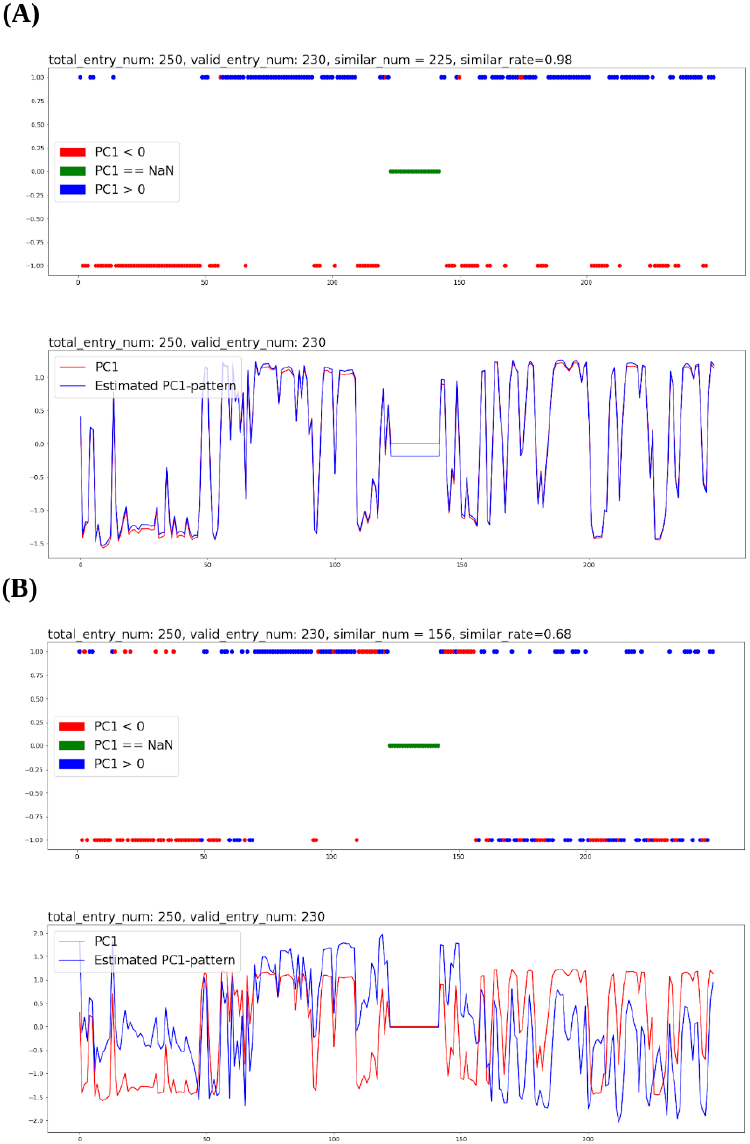
Representative comparison between the Estimated PC1-pattern and the ground-truth PC1 for K562 chromosome 1 at 1 Mb resolution. (A) Using 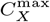 yields high agreement in both sign pattern and relative magnitude. (B) Using 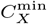 yields poor agreement. Top: sign-pattern comparison. Bottom: Z-score normalized profile comparison.

To test the generality of this observation, we evaluated all available chromosomes (excluding mtDNA) from eight GSE63525 cell lines (CH12-LX, GM12878, HMEC, HUVEC, IMR90, K562, KBM7, and NHEK) and two GSE18199 cell lines (GM06690 and K562) at 1 Mb and 100 Kb resolution. Across datasets, selecting 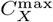 consistently produced high agreement with the ground-truth PC1, with average *similar rate* values exceeding 0.96 across chromosomes and cell lines. In contrast, selecting 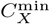 produced substantially lower agreement, typically below 0.71 on average. These results show that the dominant covariance column is a robust surrogate for PC1 in standard A/B compartment analysis. Detailed chromosome-level summaries are provided in Supplementary Table S1.

### PC1 explained variance predicts estimation reliability

We next examined whether the explained variance ratio of PC1 is associated with the reliability of HiCPEP. Across most cell lines and chromosomes, the explained variance ratio of PC1 was high at 1 Mb and 100 Kb resolution, typically exceeding 0.67 on average. Under these conditions, HiCPEP accurately recovered the PC1 sign pattern. A representative summary for K562 is shown in Table 1.

**Table 1.** Representative explained variance ratios of the first three principal components for K562 (GSE63525).

| chromosome | valid_entry_num | 1Mb |  |  | 100Kb |  |  |  |
| --- | --- | --- | --- | --- | --- | --- | --- | --- |
|  |  | Explained variance ratio |  |  | Explained variance ratio |  |  |  |
|  |  | PC1 | PC2 | PC3 | valid_entry_num | PC1 | PC2 | PC3 |
| chr1 | 230 | 0.85 | 0.08 | 0.05 | 2275 | 0.89 | 0.06 | 0.03 |
| chr2 | 242 | 0.91 | 0.05 | 0.02 | 2390 | 0.94 | 0.03 | 0.01 |
| chr3 | 196 | 0.87 | 0.05 | 0.04 | 1951 | 0.93 | 0.02 | 0.02 |
| chr4 | 190 | 0.83 | 0.14 | 0.01 | 1882 | 0.92 | 0.05 | 0.00 |
| chr5 | 179 | 0.72 | 0.24 | 0.02 | 1779 | 0.87 | 0.09 | 0.01 |
| chr6 | 170 | 0.91 | 0.05 | 0.02 | 1679 | 0.94 | 0.03 | 0.02 |
| chr7 | 158 | 0.85 | 0.07 | 0.04 | 1562 | 0.90 | 0.05 | 0.02 |
| chr8 | 145 | 0.93 | 0.03 | 0.02 | 1433 | 0.96 | 0.02 | 0.01 |
| chr9 | 125 | 0.85 | 0.10 | 0.03 | 1204 | 0.90 | 0.06 | 0.02 |
| chr10 | 134 | 0.86 | 0.05 | 0.04 | 1323 | 0.92 | 0.03 | 0.02 |
| chr11 | 133 | 0.86 | 0.08 | 0.04 | 1315 | 0.90 | 0.06 | 0.02 |
| chr12 | 132 | 0.84 | 0.09 | 0.04 | 1310 | 0.87 | 0.05 | 0.04 |
| chr13 | 97 | 0.80 | 0.10 | 0.07 | 960 | 0.83 | 0.11 | 0.04 |
| chr14 | 89 | 0.90 | 0.06 | 0.03 | 883 | 0.94 | 0.03 | 0.01 |
| chr15 | 83 | 0.77 | 0.13 | 0.05 | 822 | 0.72 | 0.19 | 0.04 |
| chr16 | 81 | 0.66 | 0.16 | 0.12 | 793 | 0.74 | 0.13 | 0.08 |
| chr17 | 80 | 0.63 | 0.24 | 0.09 | 783 | 0.73 | 0.15 | 0.06 |
| chr18 | 77 | 0.89 | 0.04 | 0.03 | 750 | 0.93 | 0.02 | 0.01 |
| chr19 | 58 | 0.62 | 0.34 | 0.02 | 562 | 0.67 | 0.27 | 0.04 |
| chr20 | 61 | 0.90 | 0.07 | 0.01 | 599 | 0.88 | 0.08 | 0.03 |
| chr21 | 38 | 0.70 | 0.28 | 0.02 | 355 | 0.82 | 0.15 | 0.01 |
| chr22 | 36 | 0.50 | 0.31 | 0.15 | 352 | 0.55 | 0.31 | 0.09 |
| chrX | 154 | 0.91 | 0.04 | 0.03 | 1520 | 0.94 | 0.01 | 0.01 |
| chrY | 11 | 0.94 | 0.06 | 0.00 | 16 | 0.94 | 0.04 | 0.01 |
| average |  | 0.81 | 0.12 | 0.04 |  | 0.86 | 0.09 | 0.03 |

Importantly, chromosomes with lower explained variance ratios were more likely to yield weaker agreement between the Estimated PC1-pattern and the ground-truth PC1. For example, K562 chromosome 22 showed a relatively low explained variance ratio at both 1 Mb and 100 Kb resolution, and correspondingly had a much smaller fraction of covariance columns that closely matched PC1. This supports our central hypothesis that the explained variance ratio of PC1 reflects how well the dominant covariance structure captures compartment organization, and therefore predicts the reliability of HiCPEP.

At the same time, the explained variance ratio should be interpreted as a qualitative rather than absolute indicator. In particular, low explained variance generally indicates weaker or noisier compartment structure, but the same threshold does not uniformly separate all reliable and unreliable chromosomes.

### Random sampling preserves accuracy

Because many covariance columns resembled PC1 when the explained variance ratio was high, we asked whether random sampling could be used to accelerate HiCPEP. For the GSE63525 datasets, we quantified the proportion of covariance columns whose sign pattern agreed with the ground-truth PC1 above predefined thresholds (Table 2). In many chromosomes, a substantial fraction of columns provided strong PC1 approximations, supporting the use of random sampling to search for 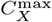 efficiently.

**Table 2.** Percentage of covariance matrix columns whose sign patterns agree with the ground-truth PC1 above predefined similarity thresholds for GSE63525 chromosomes at 1Mb and 100Kb resolution. *valid entry num* denotes the number of evaluated covariance columns. *similar rate over90, similar rate over95*, and *similar rate over99* indicate the fraction of columns whose sign pattern matches PC1 with similarity greater than 0.90, 0.95, and 0.99, respectively.

| chromosome | 1Mb |  |  |  | 100Kb |  |  |  |
| --- | --- | --- | --- | --- | --- | --- | --- | --- |
|  | valid_entry_num | similar_rate_over90 | similar_rate_over95 | similar_rate_over99 | valid_entry_num | similar_rate_over90 | similar_rate_over95 | similar_rate_over99 |
| chr1 | 230 | 0.91 | 0.81 | 0.06 | 2275 | 0.92 | 0.76 | 0.00 |
| chr2 | 242 | 0.95 | 0.85 | 0.00 | 2390 | 0.96 | 0.87 | 0.00 |
| chr3 | 196 | 0.93 | 0.40 | 0.00 | 1951 | 0.96 | 0.02 | 0.00 |
| chr4 | 190 | 0.89 | 0.73 | 0.00 | 1882 | 0.96 | 0.90 | 0.00 |
| chr5 | 179 | 0.61 | 0.34 | 0.00 | 1779 | 0.90 | 0.55 | 0.00 |
| chr6 | 170 | 0.96 | 0.91 | 0.00 | 1679 | 0.97 | 0.89 | 0.00 |
| chr7 | 158 | 0.92 | 0.58 | 0.00 | 1562 | 0.88 | 0.16 | 0.00 |
| chr8 | 145 | 0.99 | 0.97 | 0.01 | 1433 | 0.97 | 0.80 | 0.00 |
| chr9 | 125 | 0.82 | 0.63 | 0.03 | 1204 | 0.91 | 0.27 | 0.00 |
| chr10 | 134 | 0.91 | 0.78 | 0.08 | 1323 | 0.94 | 0.69 | 0.00 |
| chr11 | 133 | 0.90 | 0.75 | 0.53 | 1315 | 0.89 | 0.62 | 0.00 |
| chr12 | 132 | 0.88 | 0.22 | 0.00 | 1310 | 0.82 | 0.01 | 0.00 |
| chr13 | 97 | 0.70 | 0.09 | 0.00 | 960 | 0.83 | 0.30 | 0.00 |
| chr14 | 89 | 0.92 | 0.75 | 0.00 | 883 | 0.94 | 0.87 | 0.00 |
| chr15 | 83 | 0.75 | 0.57 | 0.00 | 822 | 0.63 | 0.42 | 0.02 |
| chr16 | 81 | 0.67 | 0.42 | 0.15 | 793 | 0.79 | 0.46 | 0.12 |
| chr17 | 80 | 0.60 | 0.41 | 0.05 | 783 | 0.79 | 0.51 | 0.02 |
| chr18 | 77 | 0.92 | 0.62 | 0.00 | 750 | 0.96 | 0.50 | 0.00 |
| chr19 | 58 | 0.07 | 0.05 | 0.03 | 562 | 0.36 | 0.30 | 0.00 |
| chr20 | 61 | 0.92 | 0.72 | 0.23 | 599 | 0.90 | 0.77 | 0.01 |
| chr21 | 38 | 0.50 | 0.42 | 0.16 | 355 | 0.87 | 0.65 | 0.33 |
| chr22 | 36 | 0.17 | 0.00 | 0.00 | 352 | 0.16 | 0.06 | 0.00 |
| chrX | 154 | 0.95 | 0.60 | 0.00 | 1520 | 0.97 | 0.38 | 0.00 |
| chrY | 11 | 1.00 | 1.00 | 1.00 | 16 | 0.94 | 0.06 | 0.06 |
| average |  | 0.79 | 0.57 | 0.10 |  | 0.84 | 0.49 | 0.02 |

We then evaluated a practical sampling strategy in which only 10% of covariance columns were examined (Table 3). On GM12878, this reduced search space still preserved high accuracy, with average *similar rate* values above 0.97 across chromosomes at 1 Mb and 100 Kb resolution, except for a small number of difficult cases such as chromosome 22 at 100 Kb resolution. These results indicate that exhaustive search over all columns is often unnecessary and that random sampling provides an effective speed–accuracy trade-off.

**Table 3.**
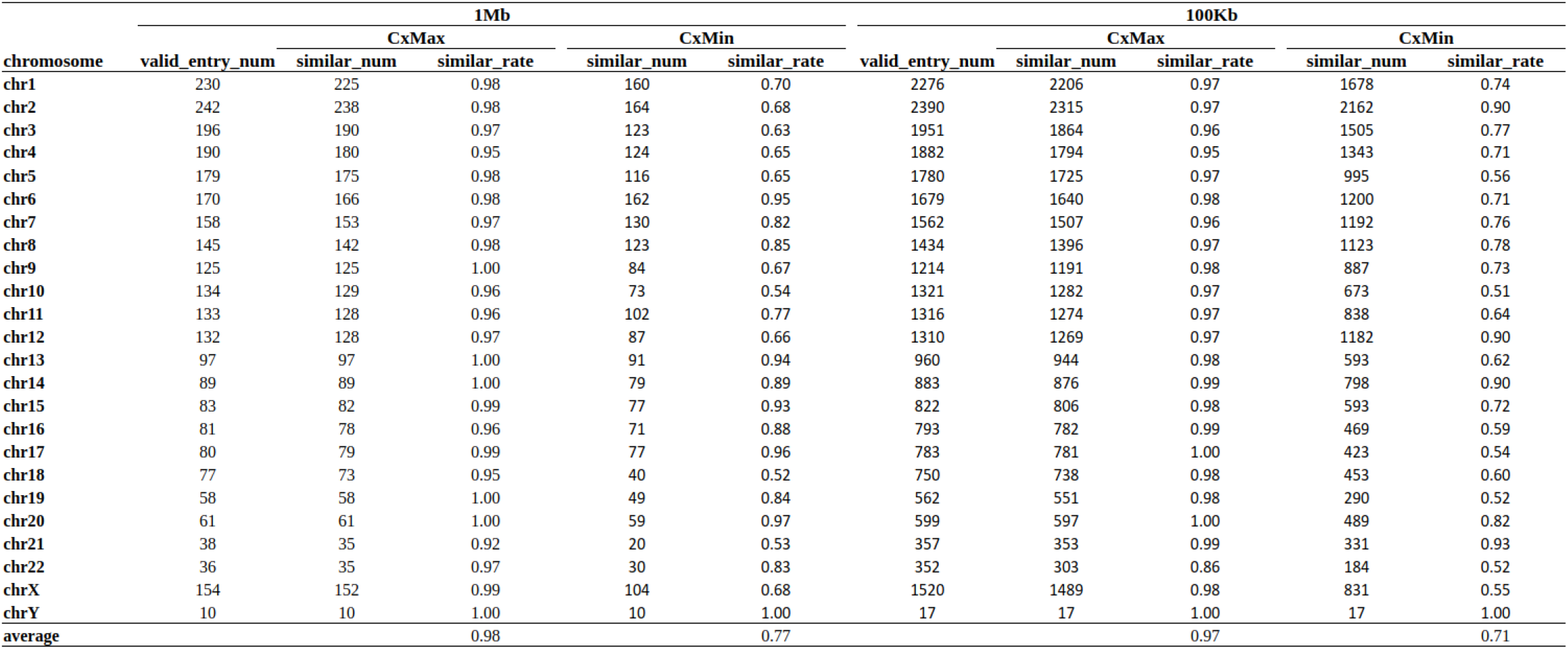
Similarity between the estimated PC1 pattern and the ground-truth PC1 for GM12878 chromosomes when only 10% of covariance columns are randomly sampled. Results are reported for 1Mb and 100Kb resolution. *similar num* and *similar rate* indicate the number and fraction of sampled columns whose sign patterns match PC1. Candidate vectors are selected using 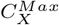 or 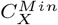.

### HiCPEP improves computational efficiency

We next compared the computational performance of HiCPEP with existing eigenvector-based approaches. In dense mode, HiCPEP was benchmarked in two settings: *Full*, which searches all covariance columns, and *Sample:0*.*1*, which evaluates only 10% of columns. On GM12878 chromosome 2, dense-mode HiCPEP achieved accuracy comparable to Scikit-learn power iteration, while using less memory and similar or slightly lower runtime (Table 4). These results show that HiCPEP can replace full eigenvector computation when a dense Pearson matrix is already available.

**Table 4.**
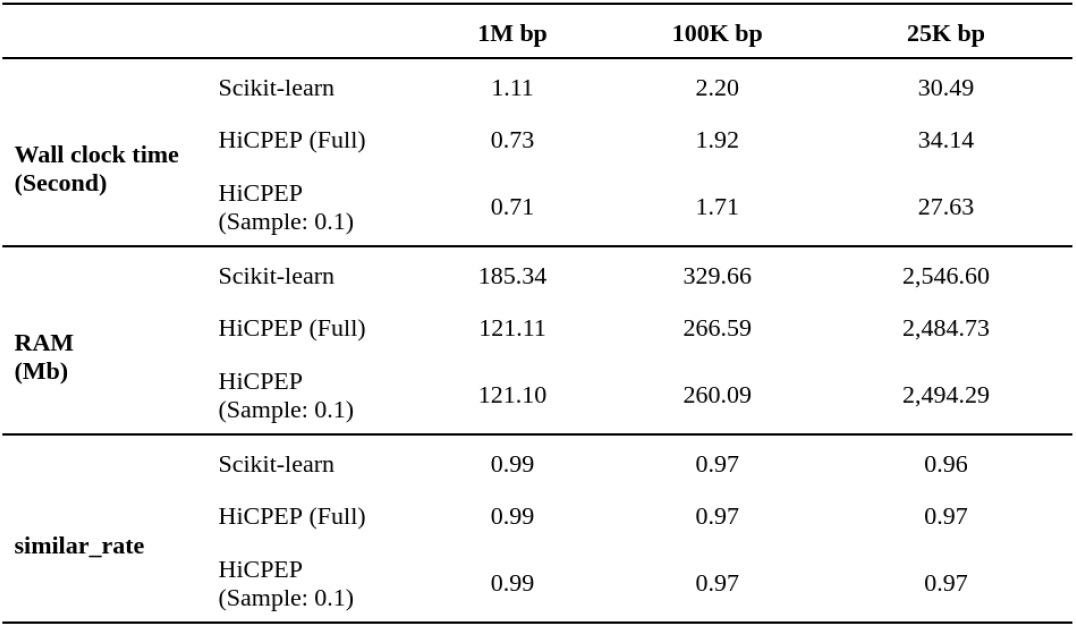
Benchmark of dense-mode HiCPEP and Scikit-learn on GM12878 chromosome 2.

|  |  | 1M bp | 100K bp | 25K bp |
| --- | --- | --- | --- | --- |
| Wall clock time<br>(Second) | Scikit-learn | 1.11 | 2.20 | 30.49 |
|  | HiCPEP (Full) | 0.73 | 1.92 | 34.14 |
|  | HiCPEP<br>(Sample: 0.1) | 0.71 | 1.71 | 27.63 |
| RAM<br>(Mb) | Scikit-learn | 185.34 | 329.66 | 2,546.60 |
|  | HiCPEP (Full) | 121.11 | 266.59 | 2,484.73 |
|  | HiCPEP<br>(Sample: 0.1) | 121.10 | 260.09 | 2,494.29 |
| similar_rate | Scikit-learn | 0.99 | 0.97 | 0.96 |
|  | HiCPEP (Full) | 0.99 | 0.97 | 0.97 |
|  | HiCPEP<br>(Sample: 0.1) | 0.99 | 0.97 | 0.97 |

We next evaluated the sparse O/E implementation of HiCPEP, which is designed to reduce memory consumption when analyzing high-resolution Hi-C data (Table 5). The current implementation is a Python prototype and is therefore not optimized for runtime; however, it demonstrates the potential memory advantages of operating directly on sparse O/E matrices. For example, at 100 Kb resolution the sparse-mode HiCPEP used substantially less RAM (146.27 MB) than dense-mode computation, while maintaining high agreement with the reference PC1 (similar rate ≥ 0.99). These results suggest that the sparse implementation is particularly suitable for high-resolution analyses or memory-constrained environments.

**Table 5.**
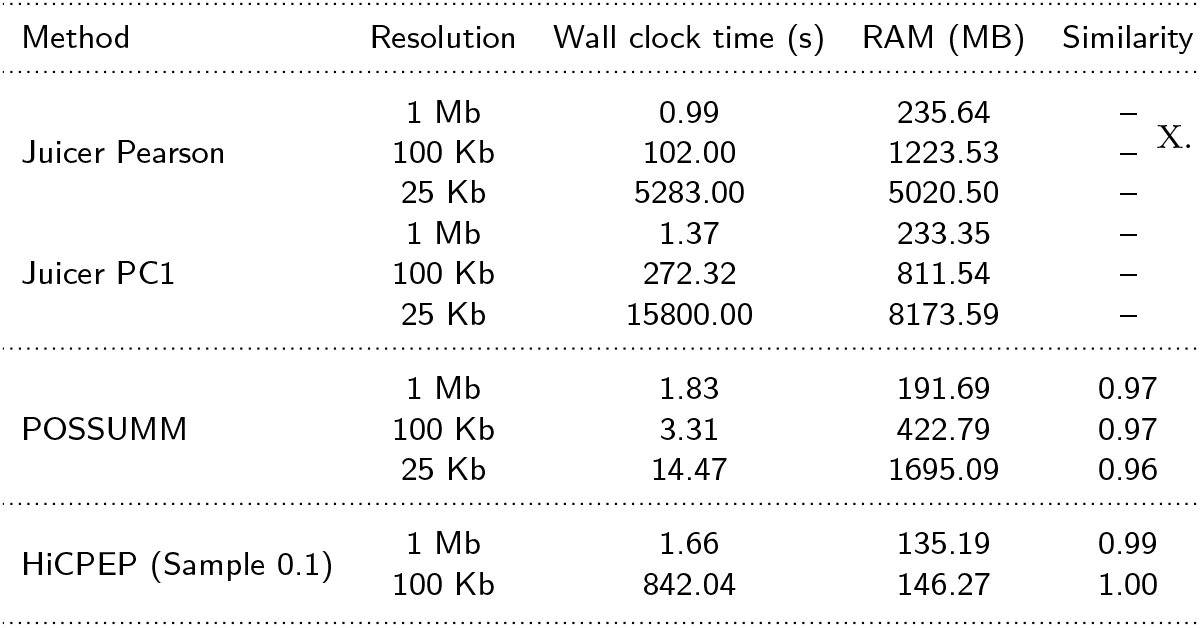
Runtime, memory usage, and similarity of PC1 estimation for GM12878 chromosome 2 across different tools and resolutions. Juicer performs the full pipeline including Pearson matrix construction and PC1 computation. POSSUMM estimates PC1 directly from Hi-C matrices. HiCPEP operates on sparse O/E matrices using 10% column sampling. The similarity score measures agreement between the estimated PC1 and the reference PC1 generated by Juicer.

| Method | Resolution | Wall clock time (s) | RAM (MB) | Similarity |
| --- | --- | --- | --- | --- |
| Juicer Pearson | 1 Mb | 0.99 | 235.64 | — |
|  | 100 Kb | 102.00 | 1223.53 | — |
|  | 25 Kb | 5283.00 | 5020.50 | — |
| Juicer PC1 | 1 Mb | 1.37 | 233.35 | — |
|  | 100 Kb | 272.32 | 811.54 | — |
|  | 25 Kb | 15800.00 | 8173.59 | — |
| POSSUMM | 1 Mb | 1.83 | 191.69 | 0.97 |
|  | 100 Kb | 3.31 | 422.79 | 0.97 |
|  | 25 Kb | 14.47 | 1695.09 | 0.96 |
| HiCPEP (Sample 0.1) | 1 Mb | 1.66 | 135.19 | 0.99 |
|  | 100 Kb | 842.04 | 146.27 | 1.00 |

We further compared HiCPEP with POSSUMM and Juicer, two widely used tools for compartment analysis. POSSUMM showed excellent runtime performance across resolutions, but required more memory than the sparse HiCPEP implementation in the tested settings (e.g., 422.79 MB vs. 146.27 MB at 100 Kb resolution). In contrast, Juicer performs the full pipeline including Pearson matrix construction and eigenvector computation, which leads to substantially higher runtime and memory consumption, particularly at fine resolutions (e.g., 15,800 s and 8,173 MB at 25 Kb resolution). Overall, these comparisons highlight complementary strengths among the methods. Juicer provides a complete end-to-end compartment analysis pipeline, POSSUMM offers highly efficient eigenvector computation from Hi-C matrices, and HiCPEP provides a practical alternative for estimating PC1 patterns either rapidly from dense Pearson matrices or with reduced memory requirements when operating directly on sparse O/E data.

## Conclusion

In this study, we investigated the relationship between the first principal component (PC1) and chromatin compartment analysis, and highlighted the role of the PC1 explained variance ratio as an indicator of compartment structure strength. Based on these observations, we proposed HiCPEP, a heuristic algorithm that approximates the sign pattern and relative magnitude of PC1 without explicit eigenvector decomposition.

HiCPEP identifies a representative column from the Hi-C Pearson covariance matrix to estimate the PC1 pattern. The method can operate either from a dense Pearson matrix for fast approximation or directly from a sparse observed/expected (O/E) matrix to reduce memory usage when analyzing high-resolution Hi-C data. In addition, because many covariance columns exhibit similar sign patterns when the explained variance ratio of PC1 is high, HiCPEP can further accelerate the estimation using random sampling without significantly sacrificing accuracy.

Across multiple Hi-C datasets, HiCPEP consistently recovered PC1 compartment patterns with high similarity to the reference PC1 generated by standard PCA-based approaches. Our experiments also show that chromosomes with higher explained variance ratios of PC1 tend to contain a larger fraction of covariance columns that are suitable for PC1 estimation, supporting the underlying intuition of the method.

Benchmark experiments demonstrate that, when the dense Pearson matrix is available, HiCPEP achieves comparable accuracy to power-iteration implementations such as Scikit-learn while requiring less memory and similar or slightly lower runtime. Furthermore, the sparse O/E implementation of HiCPEP can reduce memory usage compared with existing methods such as POSSUMM, although this memory-efficient version currently incurs higher computational cost.

Overall, HiCPEP provides a practical and scalable alternative for estimating chromatin compartment patterns from Hi-C data, bridging the gap between computational efficiency and memory constraints in large-scale compartment analysis. Future work will focus on improving the runtime of the sparse implementation and exploring integration of HiCPEP into existing Hi-C analysis pipelines.

## Data availability

All benchmarking scripts and configurations are available at https://github.com/ZhiRongDev/HiCPEP/tree/main/code_for_paper/benchmark.

The code is freely available at https://github.com/ZhiRongDev/HiCPEP.

## Author contributions statement

Z.R.C. and J.M.C. designed and conceived the study. Z.R.C. implemented the algorithm and conducted a benchmark. Z.R.C. and J.M.C. wrote and reviewed the manuscript.

## A. PCA of Cx

Set *Y* as Equation (3), where the selected *P* is a *d* × *d* orthonormal matrix that diagonalizes the covariance matrix *C*_*X*_ as Equation (4).

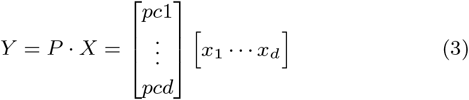

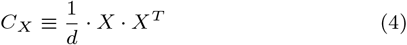

Then, the *C*_*Y*_, the derivation of PCA from Jonathon Shlens, 2014 (page 6, section V) Shlens (2014) can be adjusted as follows: The trick is to set *P* as the eigenvectors of *C*_*X*_, which implies the rows of *P* are the principal components of *X*. All the principal components are unit vectors and orthogonal to each other, and note that *C*_*X*_ is symmetric, which is orthogonally diagonalizable.

Since the principal components *P* is an orthonormal matrix (*P* ^−1^ ≡ *P*^*T*^), the covariance matrix *C*_*Y*_ can be rewritten as Equation (5), where *D* is a rank-ordered diagonal matrix from the highest to the lowest as Equation (6).

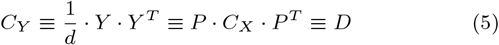

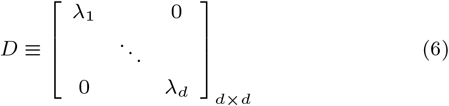

The *i*^*th*^ diagonal value of *D* (i.e., *λ*_*i*_, which is larger than or equal to 0, since *C*_*X*_ is a symmetric positive semidefinite with all real numbers.) is an eigenvalue that represents the variance of *X* along the *i*^*th*^ principal component and the definition of explained variance ratio is 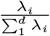. Note that the goal of PCA is to successfully find a principal component that maximizes the diagonal value of *D*, from *λ*_1_ to *λ*_*d*_, and hence the relative-magnitude of the *i*^*th*^ row of *V* will also be rank-ordered according to its multiplication factor *λ*_*i*_.

## B. Plaid pattern and the role of explained variance

Hi-C Pearson correlation matrices often exhibit a characteristic checkerboard or “plaid” pattern that reflects the segregation of genomic loci into A and B chromatin compartments. This structure is also visible in the covariance matrix derived from the Pearson matrix (Figure 2). In such cases, columns of *C*_*X*_ tend to form two groups with opposite sign patterns, corresponding to loci belonging to the A or B compartment.

**Figure 2.**
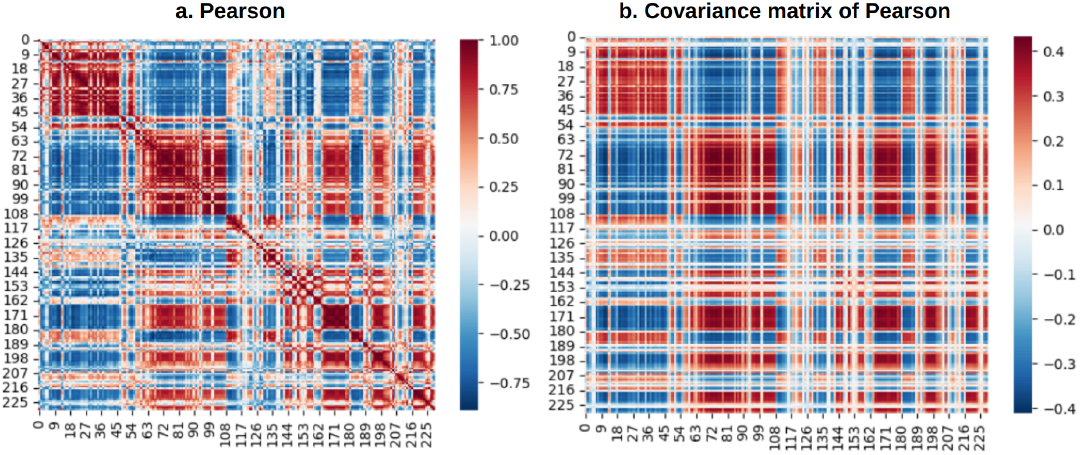
(a) Pearson correlation matrix and (b) covariance matrix of GM12878 chromosome 1 from GSE63525 at 1 Mb resolution. Both matrices display a clear plaid pattern reflecting chromatin compartment structure.

When the plaid structure is strong, the first principal component (PC1) captures the dominant variation associated with compartment organization. As a result, the explained variance ratio of the leading eigenvalue *λ*_1_ is typically high.

Figure 3 illustrates this relationship by comparing a randomly simulated Pearson matrix with a real Hi-C Pearson matrix derived from GM12878 chromosome 1 at 1 Mb resolution. The simulated matrix (Fig. 3a) does not exhibit any structured pattern and therefore produces a very small explained variance ratio for PC1 (0.029). Consequently, the PC1 track does not correspond to any meaningful genomic segregation.

**Figure 3.**
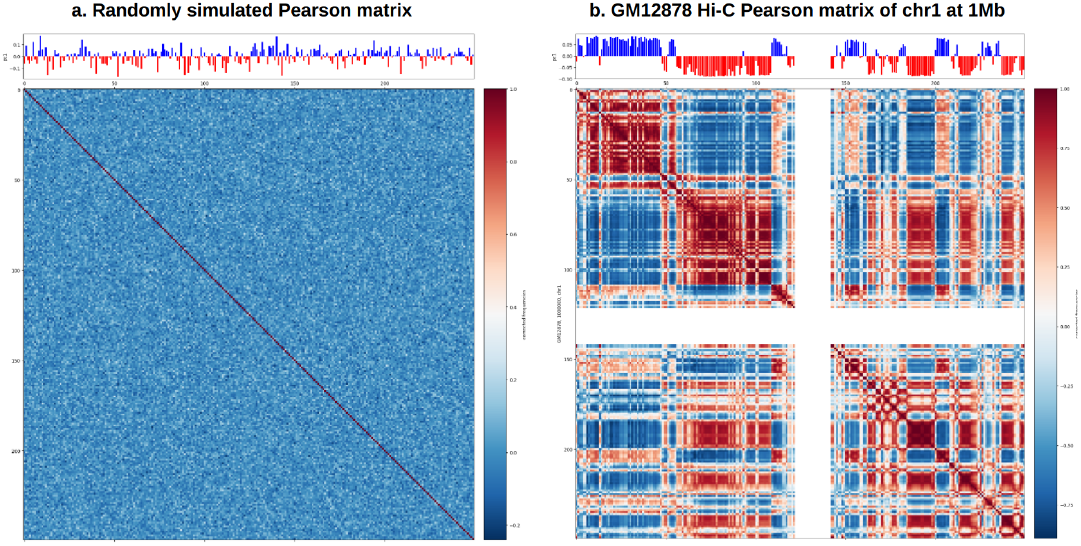
(a) Randomly simulated 250 × 250 Pearson matrix with no apparent structure and low PC1 explained variance ratio, and (b) Hi-C Pearson matrix of GM12878 chromosome 1 showing a clear plaid pattern and high PC1 explained variance ratio.

In contrast, the Hi-C Pearson matrix (Fig. 3b) shows a pronounced plaid structure. In this case, the explained variance ratio of PC1 is high (0.845), and the PC1 track clearly separates genomic regions into two opposing groups corresponding to A and B compartments.

Intuitively, when the plaid pattern is present, the PC1 vector that maximizes *λ*_1_ resembles one of the dominant sign patterns shared by many columns of *C*_*X*_. Because columns within each compartment group exhibit similar magnitudes and signs, the inner product *PC*_1_ · *C*_*X*_ yields consistently large values, resulting in a dominant eigenvalue. Conversely, when the Pearson matrix lacks a clear plaid structure, the columns of *C*_*X*_ do not align with a common sign pattern. In such cases, PC1 does not strongly resemble most covariance columns, and the explained variance ratio of *λ*_1_ remains small.

These observations suggest that the explained variance ratio of PC1 provides a useful indicator of how strongly the dominant covariance structure reflects chromatin compartment organization. A high explained variance ratio typically indicates that PC1 captures the primary compartment signal, whereas a low ratio suggests weaker or noisier compartment structure.

## C. Mathematical rationale of HiCPEP

### PCA notation in terms of the Hi-C Pearson covariance matrix

Let *X* be the *d* × *d* centered Pearson correlation matrix derived from Hi-C data, and let *C*_*X*_ denote its covariance matrix. PCA diagonalizes *C*_*X*_ as

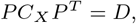

where *P* contains the eigenvectors (principal components) and *D* = diag(*λ*_1_, …, *λ*_*d*_) contains eigenvalues in descending order.

Define

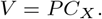

Because *pc*_*i*_ is an eigenvector of *C*_*X*_, we have

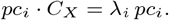

Therefore, the *i*th row of *V* equals

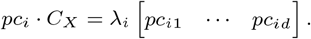

This identity shows that the entire row is the principal component scaled by its eigenvalue. Two consequences are relevant to HiCPEP:

1. **Row magnitudes reflect eigenvalues**. Since each *pc*_*i*_ is a unit vector, the relative magnitude of the *i*th row of *V* is governed by *λ*_*i*_. Thus, the first row has the largest magnitude when *λ*_1_ dominates the spectrum.
2. **Explained variance depends on alignment with** *C*_*X*_. The value of *λ*_*i*_ reflects how strongly *pc*_*i*_ aligns with the dominant column patterns in *C*_*X*_. In particular, a large *λ*_1_ indicates that *pc*_1_ matches the dominant plaid-structured variation in the Hi-C covariance matrix.

Figure 4 provides a visual interpretation of this identity. Although *V* appears to be composed of many separate multiplications, every entry in the *i*th row of *V* is simply a scaled copy of *pc*_*i*_, with scaling factor *λ*_*i*_. When *pc*_1_ aligns strongly with the major column patterns in *C*_*X*_, the first row of *V* contains large entries in absolute value, producing a large *λ*_1_.

**Figure 4.**
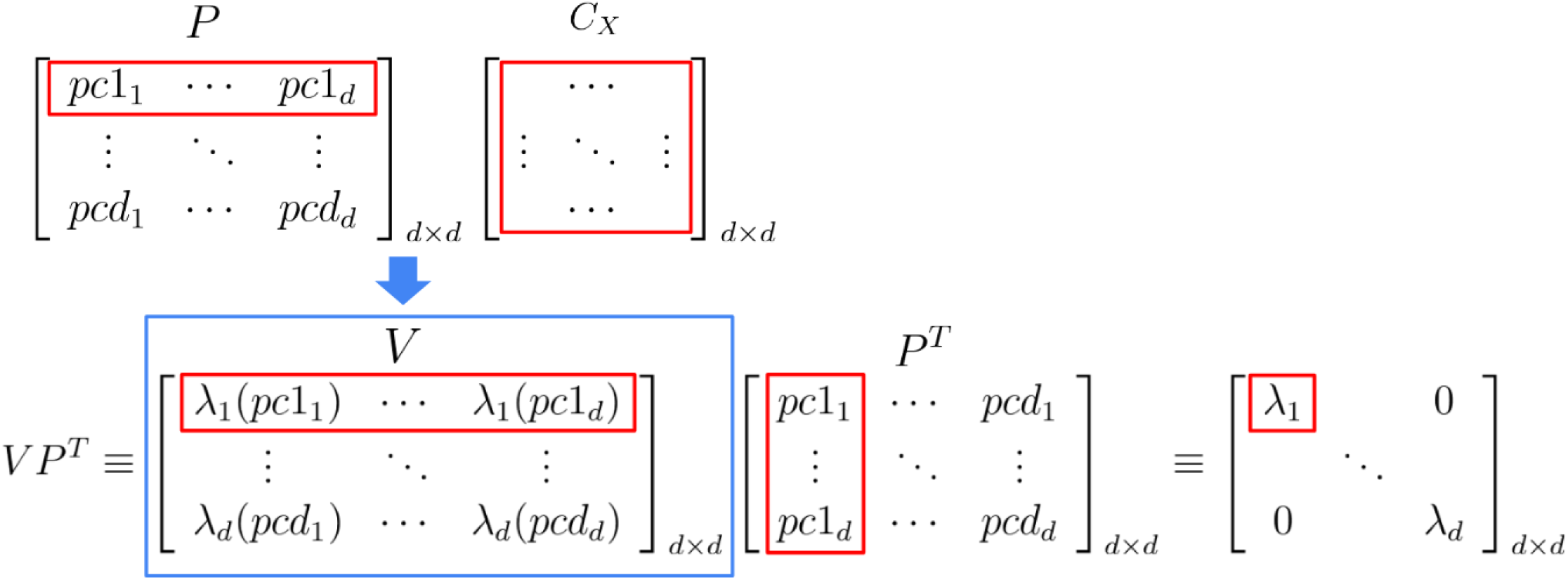
This figure shows how *λ*_1_ in *D* is determined. The red boxes indicate the rows and columns contributing to the calculation of *λ*_1_, and the blue box highlights that *V* is the matrix product of *P* and *C*_*X*_. The value of *λ*_1_ reflects the relative magnitude of the first row of *V*, which depends on how well *pc*_1_ matches the columns of *C*_*X*_.

### Why selecting 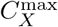 approximates pc_1_

Based on the observations above, when the explained variance ratio of *λ*_1_ is high, the sign pattern and relative magnitudes of *pc*_1_ should align with the dominant column patterns of *C*_*X*_. Because the first row of *V* = *PC*_*X*_ equals *λ*_1_*pc*_1_, the entries in this row become large precisely when *pc*_1_ matches the major column structure of *C*_*X*_. This motivates searching for a single representative column of *C*_*X*_.

We define

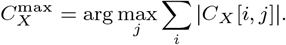

The intuition is that the column with the largest absolute column sum captures the strongest relative-magnitude pattern in *C*_*X*_ (Figure 5). When *λ*_1_ is large, this column should exhibit a sign pattern and magnitude profile similar to *pc*_1_, up to a global sign flip.

**Figure 5.**
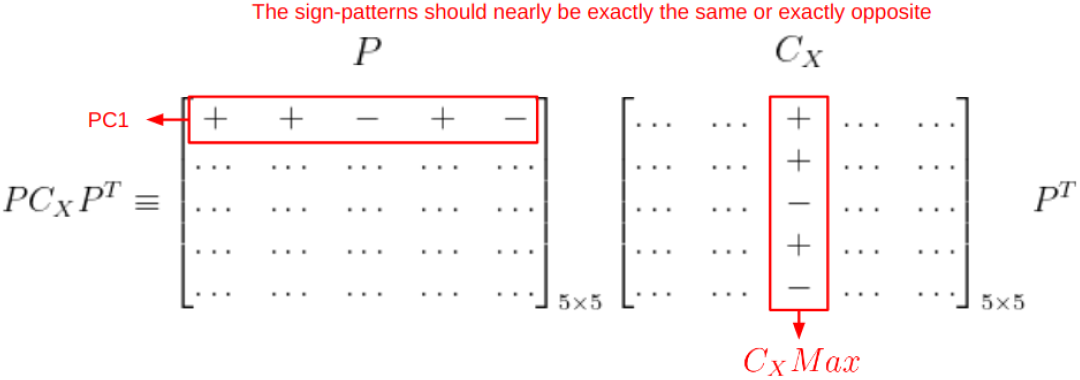
If the explained variance ratio of *λ*_1_ is high, the sign pattern and relative magnitude of *pc*_1_ can be estimated by selecting 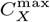 in *C*_*X*_ as the Estimated PC1-pattern.

To illustrate this intuition, we applied PCA to a simulated 5 × 5 Pearson matrix (Figure 6).

**Figure 6.**
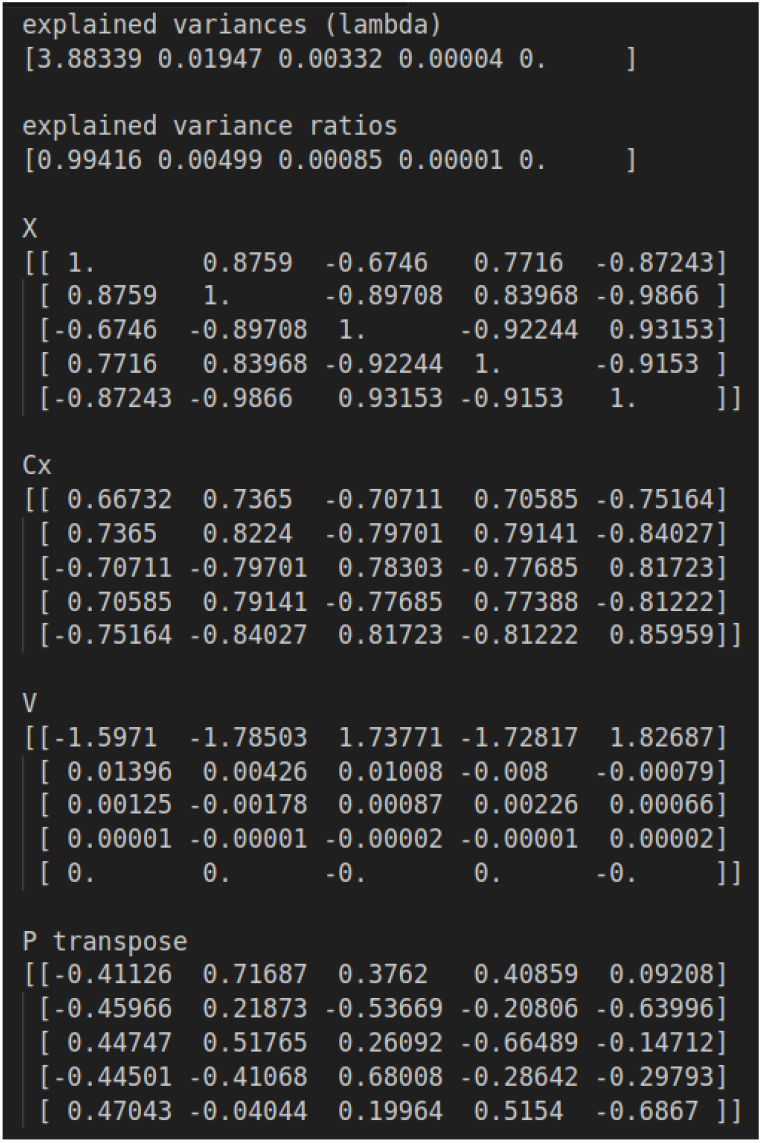
Summary of PCA on a simulated 5 × 5 Pearson matrix. The magnitudes of rows in *V* follow the order of eigenvalues.

In this example, when the explained variance ratio exceeds 0.99, the sign pattern and relative magnitudes of *C*^max^ almost perfectly reproduce those of *pc*_1_ (Figure 7). This observation is consistent with the behavior of real Hi-C Pearson matrices that exhibit strong plaid patterns.

**Figure 7.**
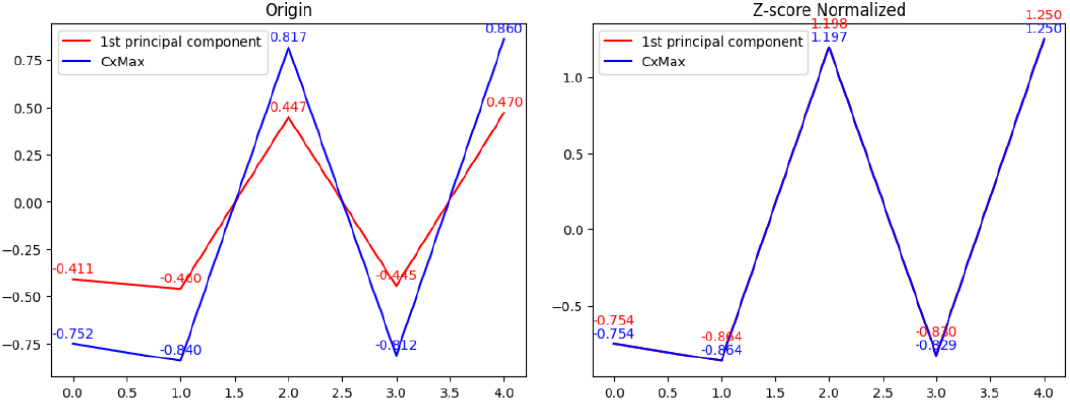
Comparison between 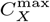 (blue) and *pc*_1_ (red). When the explained variance ratio is high, the Z-score normalized curves nearly overlap.

**Figure 8.**
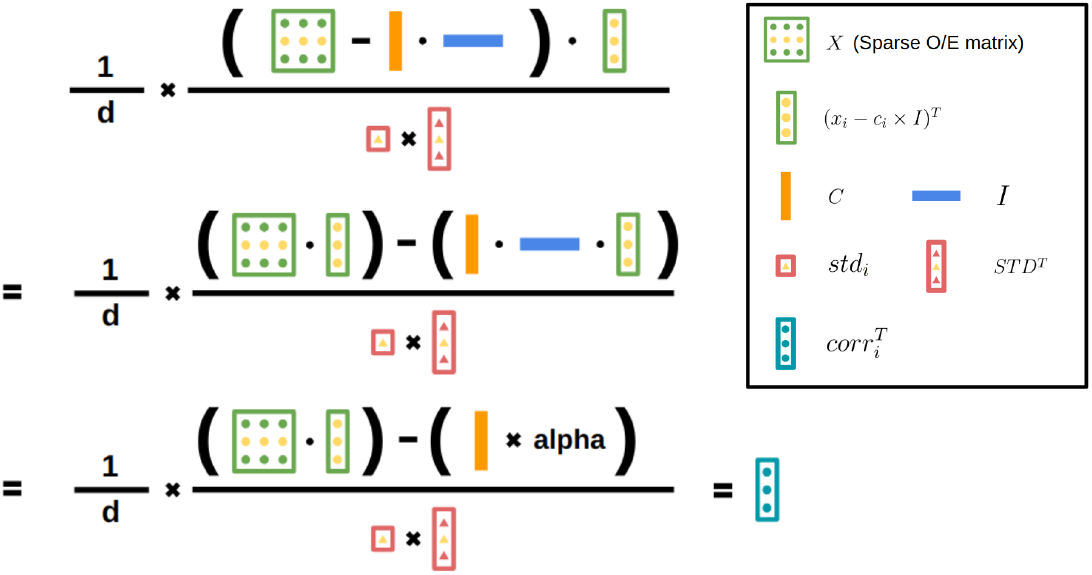
Illustration of Equation (8). The yellow dots indicate the selected row or entry used to compute *corr*_*i*_.

**Figure 9.**
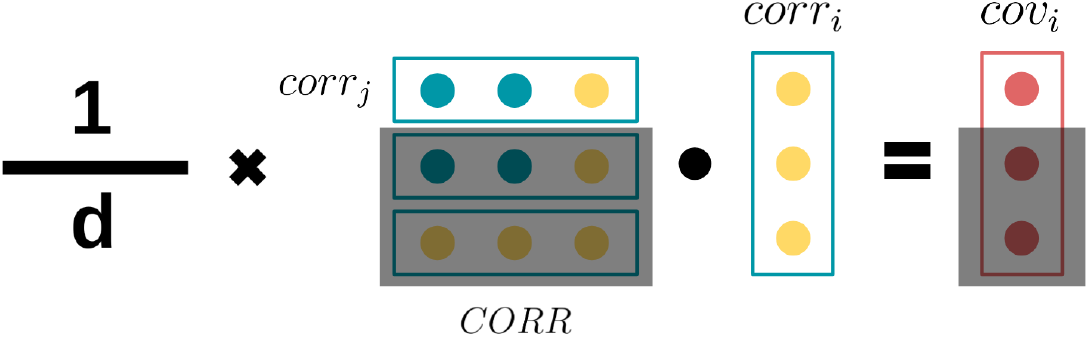
The vector *cov*_*i*_ represents the *i*th column of the Hi-C Pearson covariance matrix. Each entry is computed from the centered *corr*_*i*_ and *corr*_*j*_, generated on demand.

## D. Sparse O/E derivation

### Notation

Let the sparse O/E matrix *X* be

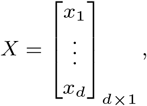

where each *x*_*i*_ is a 1 × *d* row vector. Define

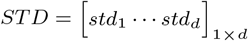

and

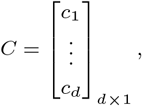

where *std*_*i*_ and *c*_*i*_ denote the standard deviation and mean of row *x*_*i*_, respectively. Let

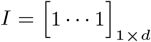

be the all-ones row vector.

The Pearson correlation matrix is written as

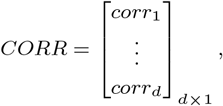

where each *corr*_*i*_ is a 1 × *d* row vector. Let

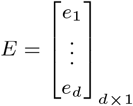

collect the row means of *CORR*, so that *e*_*i*_ is the mean of *corr*_*i*_. We assume that the sparse O/E matrix *X* can be stored in memory, but the dense Pearson matrix cannot.

### Reconstructing Pearson rows from the sparse O/E matrix

The full dense Pearson matrix can be written as

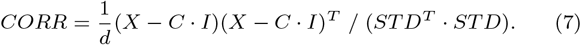

Instead of explicitly forming *CORR*, we compute one row or column at a time. For the *i*th row,

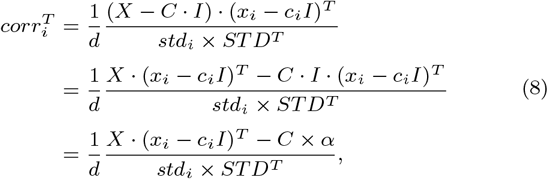

where *α* is a scalar.

Equation (8) replaces the need to store the full dense Pearson matrix with sparse matrix–vector products and *O*(*d*) intermediate vectors.

### Constructing covariance columns without a dense Pearson matrix

The covariance matrix of the Hi-C Pearson matrix is

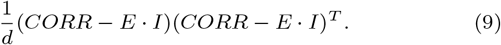

Using Eq. (8), any column of the covariance matrix can be constructed sequentially.

To compute the *i*th column *cov*_*i*_:

1. Compute *corr*_*i*_ using Eq. (8) and center it: *corr*_*i*_ ← *corr*_*i*_ − *e*_*i*_ · *I*.
2. For each *j* = 1, …, *d*:
  a. Compute *corr*_*j*_ using Eq. (8) and center it: *corr*_*j*_ ← *corr*_*j*_ *− e*_*j*_ *· I*.
  b. Set

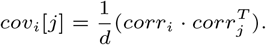

Repeating this procedure for each candidate column and tracking the column with the largest absolute sum yields *C*^max^, i.e., the Estimated PC1-pattern.

The corresponding pseudocode is shown below.

#### Algorithm 3

HiCPEP from a sparse O/E matrix (detailed form)

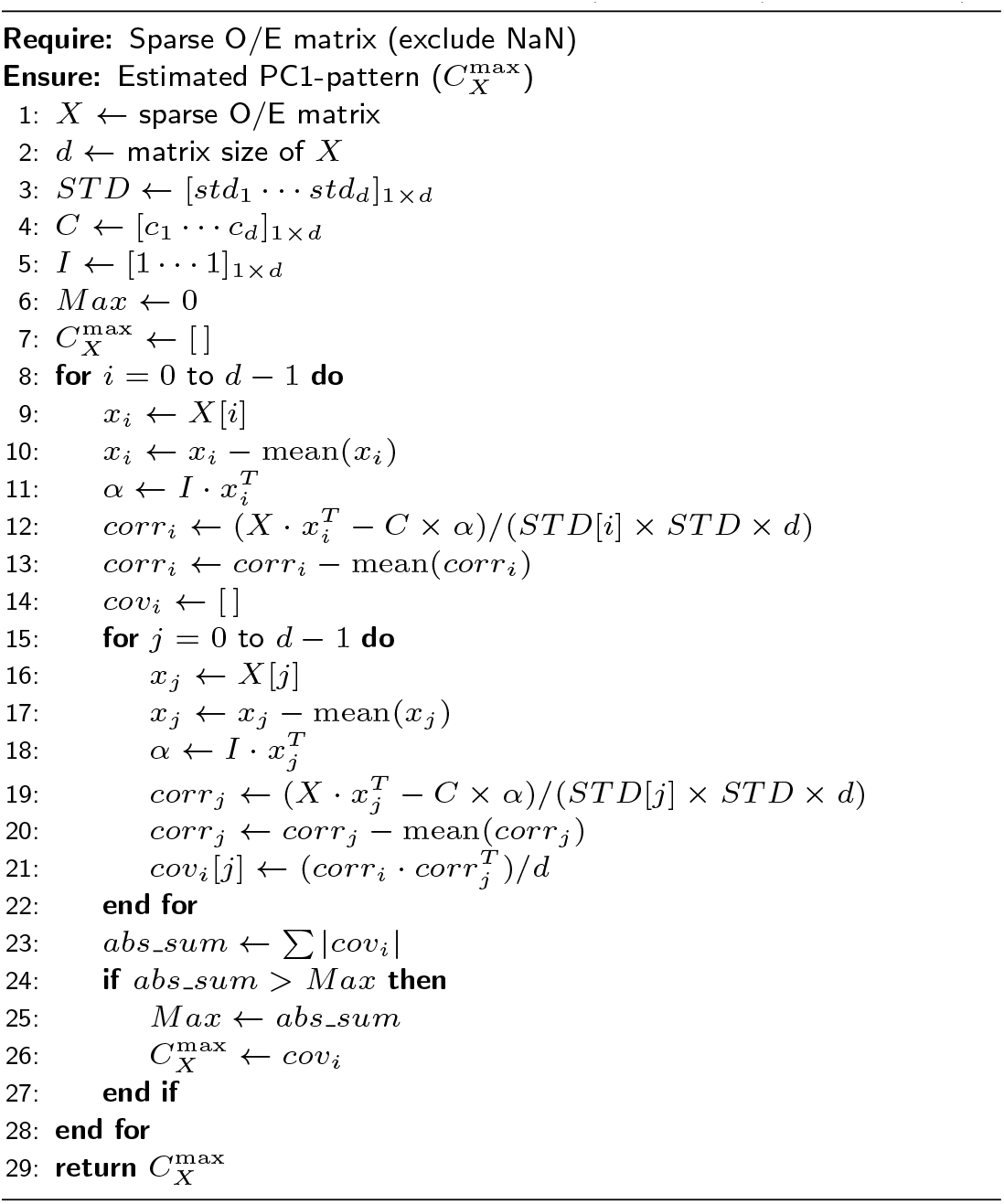

## Supplementary Tables

**Table 1.**
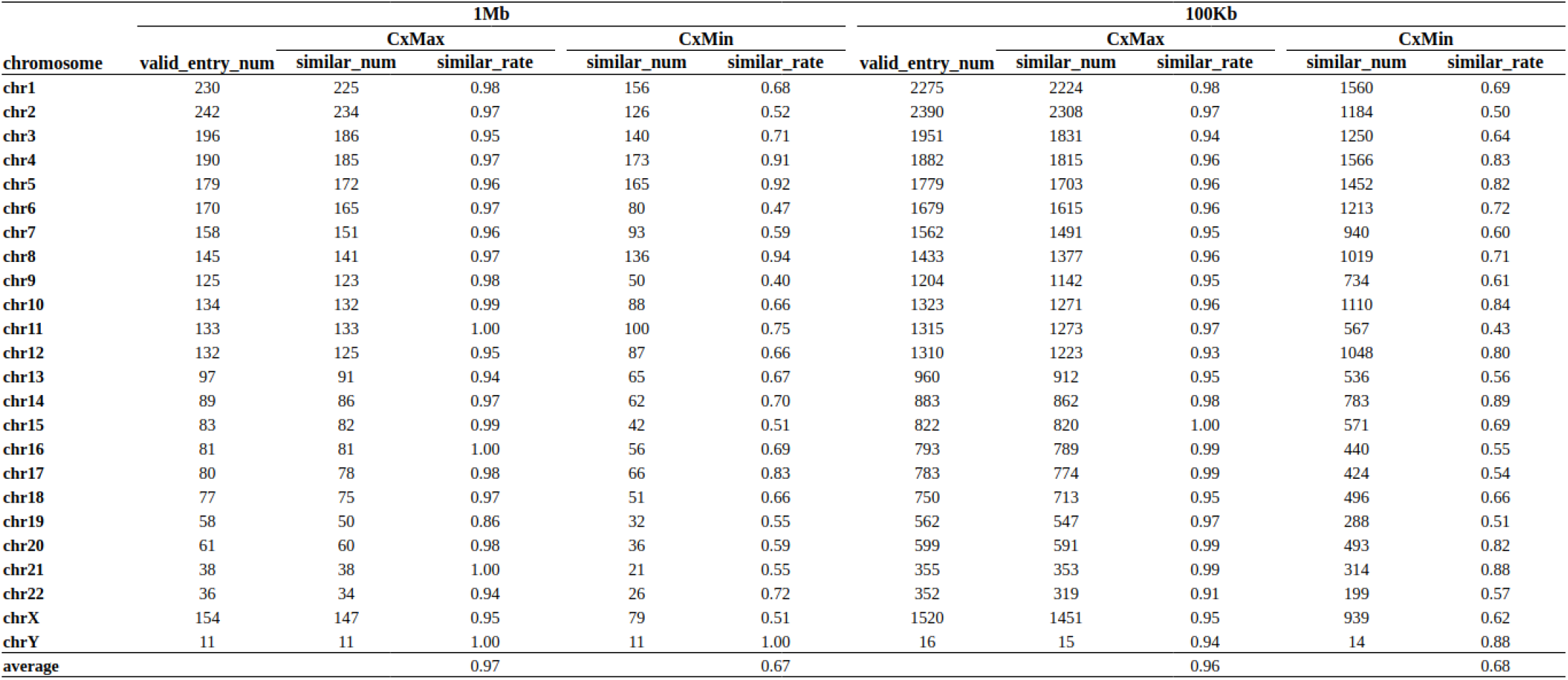
GSE63525 K562 summary for similarity (All the float numbers are rounded to the second decimal).

