## supplemental data for "HiCPEP: Efficient estimation of chromatin compartment PC1 from Hi-C covariance structure"

#### A. PCA of Cx

Set $Y$ as Equation (3), where the selected $P$ is a $d \times d$ orthonormal matrix that diagonalizes the covariance matrix $C_X$ as Equation (4).

$$Y = P \cdot X = \begin{bmatrix} pc1 \\ \vdots \\ pcd \end{bmatrix} \begin{bmatrix} x_1 \cdots x_d \end{bmatrix} \tag{3}$$

$$C_X \equiv \frac{1}{d} \cdot X \cdot X^T \tag{4}$$

Then, the $C_Y$, the derivation of PCA from Jonathon Shlens, 2014 (page 6, section V) Shlens (2014) can be adjusted as follows: The trick is to set $P$ as the eigenvectors of $C_X$, which implies the rows of $P$ are the principal components of $X$. All the principal components are unit vectors and orthogonal to each other, and note that $C_X$ is symmetric, which is orthogonally diagonalizable.

Since the principal components $P$ is an orthonormal matrix ($P^{-1} \equiv P^T$), the covariance matrix $C_Y$ can be rewritten as Equation (5), where $D$ is a rank-ordered diagonal matrix from the highest to the lowest as Equation (6).

$$C_Y \equiv \frac{1}{d} \cdot Y \cdot Y^T \equiv P \cdot C_X \cdot P^T \equiv D \tag{5}$$

$$D \equiv \begin{bmatrix} \lambda_1 & & 0 \\ & \ddots & \\ 0 & & \lambda_d \end{bmatrix}_{d \times d} \tag{6}$$

The $i^{th}$ diagonal value of $D$ (i.e., $\lambda_i$, which is larger than or equal to 0, since $C_X$ is a symmetric positive semidefinite with all real numbers.) is an eigenvalue that represents the variance of $X$ along the $i^{th}$ principal component and the definition of explained variance ratio is $\frac{\lambda_i}{\sum_1^d \lambda_i}$. Note that the goal of PCA is to successfully find a principal component that maximizes the diagonal value of $D$, from $\lambda_1$ to $\lambda_d$, and hence the relative-magnitude of the $i^{th}$ row of $V$ will also be rank-ordered according to its multiplication factor $\lambda_i$.

#### B. Plaid pattern and the role of explained variance

Hi-C Pearson correlation matrices often exhibit a characteristic checkerboard or “plaid” pattern that reflects the segregation of genomic loci into A and B chromatin compartments. This structure is also visible in the covariance matrix derived from the Pearson matrix (Figure 2). In such cases, columns of $C_X$ tend to form two groups with opposite sign patterns, corresponding to loci belonging to the A or B compartment.

When the plaid structure is strong, the first principal component (PC1) captures the dominant variation associated with compartment organization. As a result, the explained variance ratio of the leading eigenvalue $\lambda_1$ is typically high.

Figure 3 illustrates this relationship by comparing a randomly simulated Pearson matrix with a real Hi-C Pearson matrix derived from GM12878 chromosome 1 at 1 Mb resolution. The simulated matrix (Fig. 3a) does not exhibit any structured pattern and therefore produces a very small explained variance ratio for PC1 (0.029). Consequently, the PC1 track does not correspond to any meaningful genomic segregation.

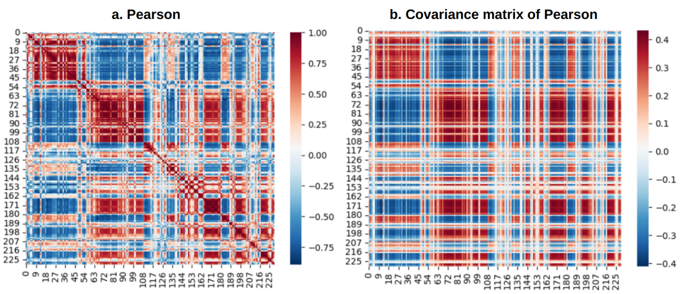


**Figure 2** (a) Pearson correlation matrix and (b) covariance matrix of GM12878 chromosome 1 from GSE63525 at 1 Mb resolution. Both matrices display a clear plaid pattern reflecting chromatin compartment structure.

In contrast, the Hi-C Pearson matrix (Fig. 3b) shows a pronounced plaid structure. In this case, the explained variance ratio of PC1 is high (0.845), and the PC1 track clearly separates genomic regions into two opposing groups corresponding to A and B compartments.

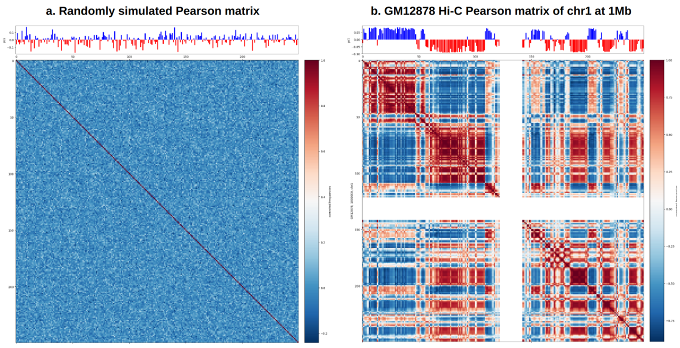


**Figure 3** (a) Randomly simulated $250 \times 250$ Pearson matrix with no apparent structure and low PC1 explained variance ratio, and (b) Hi-C Pearson matrix of GM12878 chromosome 1 showing a clear plaid pattern and high PC1 explained variance ratio.

Intuitively, when the plaid pattern is present, the PC1 vector that maximizes $\lambda_1$ resembles one of the dominant sign patterns shared by many columns of $C_X$. Because columns within each compartment group exhibit similar magnitudes and signs, the inner product $PC_1 \cdot C_X$ yields consistently large values, resulting in a dominant eigenvalue. Conversely, when the Pearson matrix lacks a clear plaid structure, the columns of $C_X$ do not align with a common sign pattern. In such cases, PC1 does not strongly resemble most covariance columns, and the explained variance ratio of $\lambda_1$ remains small.

These observations suggest that the explained variance ratio of PC1 provides a useful indicator of how strongly the dominant covariance structure reflects chromatin compartment organization. A high explained variance ratio typically indicates that PC1 captures the primary compartment signal, whereas a low ratio suggests weaker or noisier compartment structure.

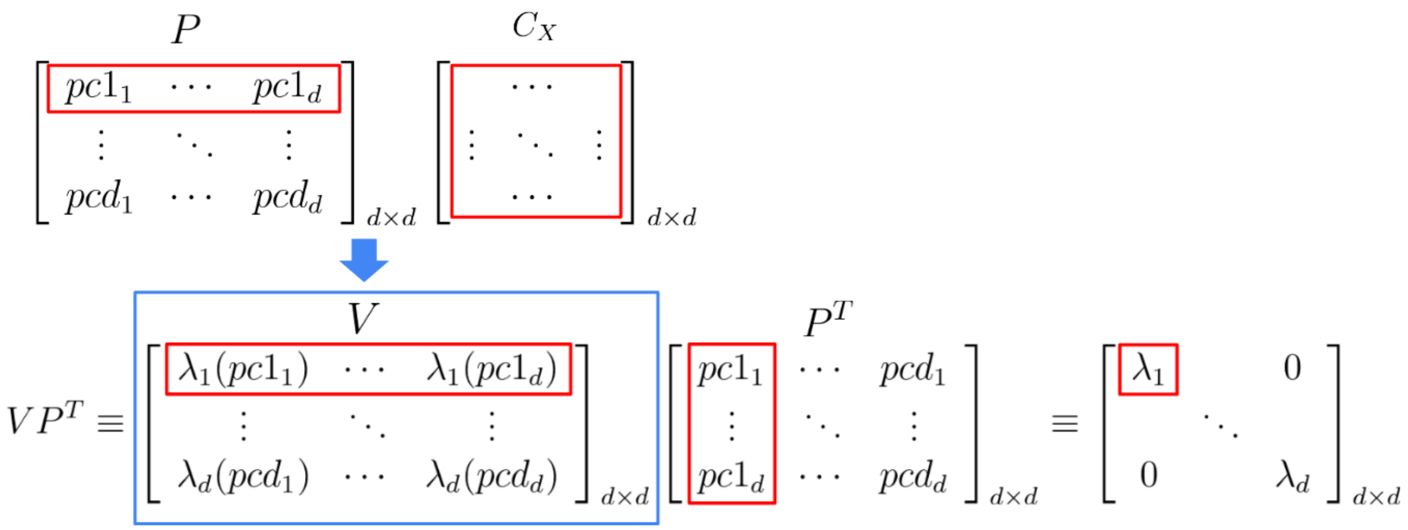


**Figure 4** This figure shows how $\lambda_1$ in $D$ is determined. The red boxes indicate the rows and columns contributing to the calculation of $\lambda_1$, and the blue box highlights that $V$ is the matrix product of $P$ and $C_X$. The value of $\lambda_1$ reflects the relative magnitude of the first row of $V$, which depends on how well $pc_1$ matches the columns of $C_X$.

#### C. Mathematical rationale of HiCPEP

##### PCA notation in terms of the Hi-C Pearson covariance matrix

Let $X$ be the $d \times d$ centered Pearson correlation matrix derived from Hi-C data, and let $C_X$ denote its covariance matrix. PCA diagonalizes $C_X$ as

$$PC_XP^T = D,$$

where $P$ contains the eigenvectors (principal components) and $D = \mathrm{diag}(\lambda_1, \dots, \lambda_d)$ contains eigenvalues in descending order.

Define

$$V = PC_X.$$

Because $pc_i$ is an eigenvector of $C_X$, we have

$$pc_i \cdot C_X = \lambda_i \, pc_i.$$

Therefore, the $i$th row of $V$ equals

$$pc_i \cdot C_X = \lambda_i \begin{bmatrix} pc_{i1} & \cdots & pc_{id} \end{bmatrix}.$$

This identity shows that the entire row is the principal component scaled by its eigenvalue.

Two consequences are relevant to HiCPEP:

1. **Row magnitudes reflect eigenvalues.** Since each $pc_i$ is a unit vector, the relative magnitude of the $i$th row of $V$ is governed by $\lambda_i$. Thus, the first row has the largest magnitude when $\lambda_1$ dominates the spectrum.
2. **Explained variance depends on alignment with $C_X$.** The value of $\lambda_i$ reflects how strongly $pc_i$ aligns with the dominant column patterns in $C_X$. In particular, a large $\lambda_1$ indicates that $pc_1$ matches the dominant plaid-structured variation in the Hi-C covariance matrix.

Figure 4 provides a visual interpretation of this identity. Although $V$ appears to be composed of many separate multiplications, every entry in the $i$th row of $V$ is simply a scaled copy of $pc_i$, with scaling factor $\lambda_i$. When $pc_1$ aligns strongly with the major column patterns in $C_X$, the first row of $V$ contains large entries in absolute value, producing a large $\lambda_1$.

##### Why selecting $C_X^{\max}$ approximates $pc_1$

Based on the observations above, when the explained variance ratio of $\lambda_1$ is high, the sign pattern and relative magnitudes of $pc_1$ should align with the dominant column patterns of $C_X$. Because the first row of $V = PC_X$ equals $\lambda_1 pc_1$, the entries in this row become large precisely when $pc_1$ matches the major column structure of $C_X$. This motivates searching for a single representative column of $C_X$.

We define

$$C_X^{\max} = \arg\max_j \sum_i |C_X[i,j]|.$$

The intuition is that the column with the largest absolute column sum captures the strongest relative-magnitude pattern in $C_X$ (Figure 5). When $\lambda_1$ is large, this column should exhibit a sign pattern and magnitude profile similar to $pc_1$, up to a global sign flip.

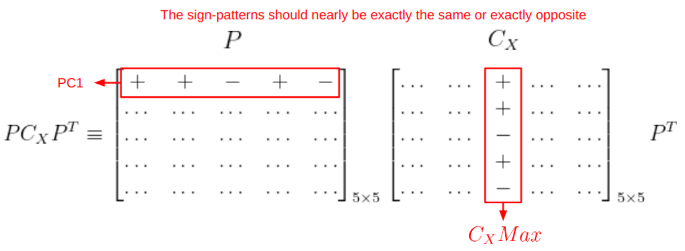


**Figure 5** If the explained variance ratio of $\lambda_1$ is high, the sign pattern and relative magnitude of $pc_1$ can be estimated by selecting $C_X^{\max}$ in $C_X$ as the Estimated PC1-pattern.

To illustrate this intuition, we applied PCA to a simulated $5 \times 5$ Pearson matrix (Figure 6).

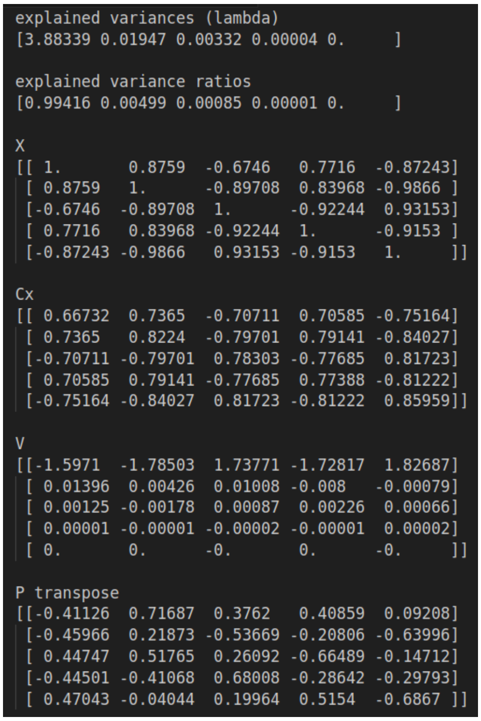

```
explained variances (lambda)
[3.88339 0.01947 0.00332 0.00004 0.     ]

explained variance ratios
[0.99416 0.00499 0.00085 0.00001 0.     ]

X
[[ 1.       0.8759  -0.6746   0.7716  -0.87243]
 [ 0.8759   1.      -0.89708  0.83968 -0.9866 ]
 [-0.6746  -0.89708  1.      -0.92244  0.93153]
 [ 0.7716   0.83968 -0.92244  1.      -0.9153 ]
 [-0.87243 -0.9866   0.93153 -0.9153   1.     ]]

Cx
[[ 0.66732  0.7365  -0.70711  0.70585 -0.75164]
 [ 0.7365   0.8224  -0.79701  0.79141 -0.84027]
 [-0.70711 -0.79701  0.78303 -0.77685  0.81723]
 [ 0.70585  0.79141 -0.77685  0.77388 -0.81222]
 [-0.75164 -0.84027  0.81723 -0.81222  0.85959]]

V
[[-1.5971  -1.78503  1.73771 -1.72817  1.82687]
 [ 0.01396  0.00426  0.01008 -0.008   -0.00079]
 [ 0.00125 -0.00178  0.00087  0.00226  0.00066]
 [ 0.00001 -0.00001 -0.00002 -0.00001  0.00002]
 [ 0.       0.      -0.       0.      -0.     ]]

P transpose
[[-0.41126  0.71687  0.3762   0.40859  0.09208]
 [-0.45966  0.21873 -0.53669 -0.20806 -0.63996]
 [ 0.44747  0.51765  0.26092 -0.66489 -0.14712]
 [-0.44501 -0.41068  0.68008 -0.28642 -0.29793]
 [ 0.47043 -0.04044  0.19964  0.5154  -0.6867 ]]
```

**Figure 6** Summary of PCA on a simulated $5 \times 5$ Pearson matrix. The magnitudes of rows in $V$ follow the order of eigenvalues.

In this example, when the explained variance ratio exceeds 0.99, the sign pattern and relative magnitudes of $C_X^{\max}$ almost perfectly reproduce those of $pc_1$ (Figure 7). This observation is consistent with the behavior of real Hi-C Pearson matrices that exhibit strong plaid patterns.

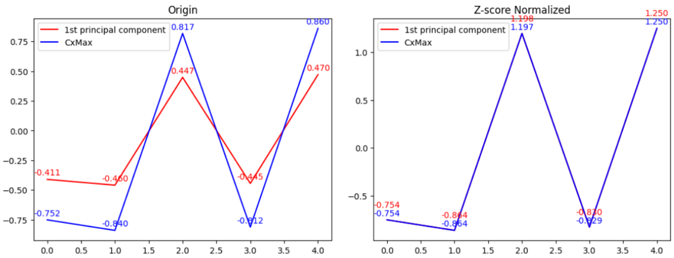


**Figure 7** Comparison between $C_X^{\max}$ (blue) and $pc_1$ (red). When the explained variance ratio is high, the Z-score normalized curves nearly overlap.

### D. Sparse O/E derivation

#### Notation

Let the sparse O/E matrix $X$ be

$$X = \begin{bmatrix} x_1 \\ \vdots \\ x_d \end{bmatrix}_{d \times 1},$$

where each $x_i$ is a $1 \times d$ row vector. Define

$$STD = \begin{bmatrix} std_1 \cdots std_d \end{bmatrix}_{1 \times d}$$

and

$$C = \begin{bmatrix} c_1 \\ \vdots \\ c_d \end{bmatrix}_{d \times 1},$$

where $std_i$ and $c_i$ denote the standard deviation and mean of row $x_i$, respectively. Let

$$I = \begin{bmatrix} 1 \cdots 1 \end{bmatrix}_{1 \times d}$$

be the all-ones row vector.

The Pearson correlation matrix is written as

$$CORR = \begin{bmatrix} corr_1 \\ \vdots \\ corr_d \end{bmatrix}_{d \times 1},$$

where each $corr_i$ is a $1 \times d$ row vector. Let

$$E = \begin{bmatrix} e_1 \\ \vdots \\ e_d \end{bmatrix}_{d \times 1}$$

collect the row means of $CORR$, so that $e_i$ is the mean of $corr_i$. We assume that the sparse O/E matrix $X$ can be stored in memory, but the dense Pearson matrix cannot.

#### Reconstructing Pearson rows from the sparse O/E matrix

The full dense Pearson matrix can be written as

$$CORR = \frac{1}{d}(X - C \cdot I)(X - C \cdot I)^T \; / \; (STD^T \cdot STD). \tag{7}$$

Instead of explicitly forming $CORR$, we compute one row or column at a time. For the $i$th row,

$$\begin{aligned} corr_i^T &= \frac{1}{d} \frac{(X - C \cdot I) \cdot (x_i - c_i I)^T}{std_i \times STD^T} \\ &= \frac{1}{d} \frac{X \cdot (x_i - c_i I)^T - C \cdot I \cdot (x_i - c_i I)^T}{std_i \times STD^T} \\ &= \frac{1}{d} \frac{X \cdot (x_i - c_i I)^T - C \times \alpha}{std_i \times STD^T}, \end{aligned} \tag{8}$$

where $\alpha$ is a scalar.

Equation (8) replaces the need to store the full dense Pearson matrix with sparse matrix–vector products and $O(d)$ intermediate vectors.

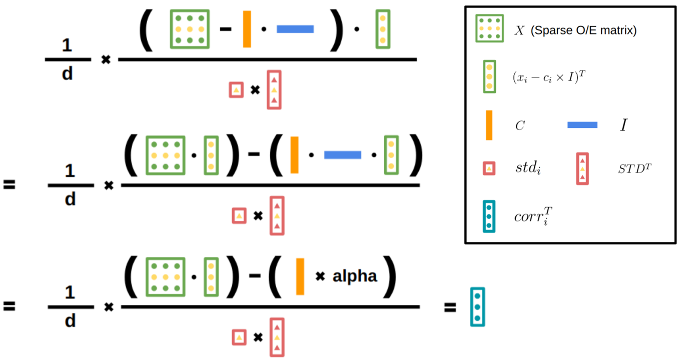


**Figure 8** Illustration of Equation (8). The yellow dots indicate the selected row or entry used to compute $corr_i$.

#### Constructing covariance columns without a dense Pearson matrix

The covariance matrix of the Hi-C Pearson matrix is

$$\frac{1}{d}(CORR - E \cdot I)(CORR - E \cdot I)^T. \quad (9)$$

Using Eq. (8), any column of the covariance matrix can be constructed sequentially.

To compute the $i$th column $cov_i$:

1. Compute $corr_i$ using Eq. (8) and center it: $corr_i \leftarrow corr_i - e_i \cdot I$.
2. For each $j = 1, \ldots, d$:
   a. Compute $corr_j$ using Eq. (8) and center it: $corr_j \leftarrow corr_j - e_j \cdot I$.
   b. Set
   $$cov_i[j] = \frac{1}{d}(corr_i \cdot corr_j^T).$$

Repeating this procedure for each candidate column and tracking the column with the largest absolute sum yields $C_X^{\max}$, i.e., the Estimated PC1-pattern.

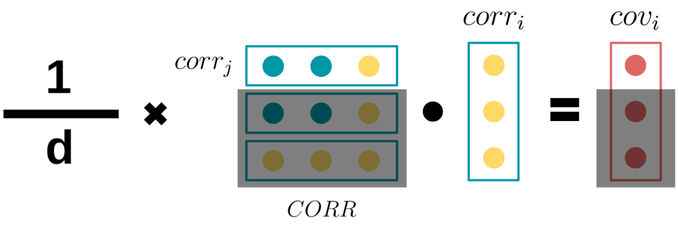


**Figure 9** The vector $cov_i$ represents the $i$th column of the Hi-C Pearson covariance matrix. Each entry is computed from the centered $corr_i$ and $corr_j$, generated on demand.

The corresponding pseudocode is shown below.

**Algorithm 3** HiCPEP from a sparse O/E matrix (detailed form)

**Require:** Sparse O/E matrix (exclude NaN)
**Ensure:** Estimated PC1-pattern ($C_X^{\max}$)

$X \leftarrow$ sparse O/E matrix
$d \leftarrow$ matrix size of $X$
$STD \leftarrow [std_1 \cdots std_d]_{1\times d}$
$C \leftarrow [c_1 \cdots c_d]_{1\times d}$
$I \leftarrow [1 \cdots 1]_{1\times d}$
$Max \leftarrow 0$
$C_X^{\max} \leftarrow [\,]$
**for** $i = 0$ to $d-1$ **do**
  $x_i \leftarrow X[i]$
  $x_i \leftarrow x_i - \text{mean}(x_i)$
  $\alpha \leftarrow I \cdot x_i^T$
  $corr_i \leftarrow (X \cdot x_i^T - C \times \alpha)/(STD[i] \times STD \times d)$
  $corr_i \leftarrow corr_i - \text{mean}(corr_i)$
  $cov_i \leftarrow [\,]$
  **for** $j = 0$ to $d-1$ **do**
    $x_j \leftarrow X[j]$
    $x_j \leftarrow x_j - \text{mean}(x_j)$
    $\alpha \leftarrow I \cdot x_j^T$
    $corr_j \leftarrow (X \cdot x_j^T - C \times \alpha)/(STD[j] \times STD \times d)$
    $corr_j \leftarrow corr_j - \text{mean}(corr_j)$
    $cov_i[j] \leftarrow (corr_i \cdot corr_j^T)/d$
  **end for**
  $abs\_sum \leftarrow \sum |cov_i|$
  **if** $abs\_sum > Max$ **then**
    $Max \leftarrow abs\_sum$
    $C_X^{\max} \leftarrow cov_i$
  **end if**
**end for**
**return** $C_X^{\max}$

#### Supplementary Tables

**Table 1** GSE63525 K562 summary for similarity (All the float numbers are rounded to the second decimal).

| | 1Mb | | | | | 100Kb | | | | |
|---|---|---|---|---|---|---|---|---|---|---|
| | | CxMax | | CxMin | | | CxMax | | CxMin | |
| **chromosome** | **valid_entry_num** | **similar_num** | **similar_rate** | **similar_num** | **similar_rate** | **valid_entry_num** | **similar_num** | **similar_rate** | **similar_num** | **similar_rate** |
| **chr1** | 230 | 225 | 0.98 | 156 | 0.68 | 2275 | 2224 | 0.98 | 1560 | 0.69 |
| **chr2** | 242 | 234 | 0.97 | 126 | 0.52 | 2390 | 2308 | 0.97 | 1184 | 0.50 |
| **chr3** | 196 | 186 | 0.95 | 140 | 0.71 | 1951 | 1831 | 0.94 | 1250 | 0.64 |
| **chr4** | 190 | 185 | 0.97 | 173 | 0.91 | 1882 | 1815 | 0.96 | 1566 | 0.83 |
| **chr5** | 179 | 172 | 0.96 | 165 | 0.92 | 1779 | 1703 | 0.96 | 1452 | 0.82 |
| **chr6** | 170 | 165 | 0.97 | 80 | 0.47 | 1679 | 1615 | 0.96 | 1213 | 0.72 |
| **chr7** | 158 | 151 | 0.96 | 93 | 0.59 | 1562 | 1491 | 0.95 | 940 | 0.60 |
| **chr8** | 145 | 141 | 0.97 | 136 | 0.94 | 1433 | 1377 | 0.96 | 1019 | 0.71 |
| **chr9** | 125 | 123 | 0.98 | 50 | 0.40 | 1204 | 1142 | 0.95 | 734 | 0.61 |
| **chr10** | 134 | 132 | 0.99 | 88 | 0.66 | 1323 | 1271 | 0.96 | 1110 | 0.84 |
| **chr11** | 133 | 133 | 1.00 | 100 | 0.75 | 1315 | 1273 | 0.97 | 567 | 0.43 |
| **chr12** | 132 | 125 | 0.95 | 87 | 0.66 | 1310 | 1223 | 0.93 | 1048 | 0.80 |
| **chr13** | 97 | 91 | 0.94 | 65 | 0.67 | 960 | 912 | 0.95 | 536 | 0.56 |
| **chr14** | 89 | 86 | 0.97 | 62 | 0.70 | 883 | 862 | 0.98 | 783 | 0.89 |
| **chr15** | 83 | 82 | 0.99 | 42 | 0.51 | 822 | 820 | 1.00 | 571 | 0.69 |
| **chr16** | 81 | 81 | 1.00 | 56 | 0.69 | 793 | 789 | 0.99 | 440 | 0.55 |
| **chr17** | 80 | 78 | 0.98 | 66 | 0.83 | 783 | 774 | 0.99 | 424 | 0.54 |
| **chr18** | 77 | 75 | 0.97 | 51 | 0.66 | 750 | 713 | 0.95 | 496 | 0.66 |
| **chr19** | 58 | 50 | 0.86 | 32 | 0.55 | 562 | 547 | 0.97 | 288 | 0.51 |
| **chr20** | 61 | 60 | 0.98 | 36 | 0.59 | 599 | 591 | 0.99 | 493 | 0.82 |
| **chr21** | 38 | 38 | 1.00 | 21 | 0.55 | 355 | 353 | 0.99 | 314 | 0.88 |
| **chr22** | 36 | 34 | 0.94 | 26 | 0.72 | 352 | 319 | 0.91 | 199 | 0.57 |
| **chrX** | 154 | 147 | 0.95 | 79 | 0.51 | 1520 | 1451 | 0.95 | 939 | 0.62 |
| **chrY** | 11 | 11 | 1.00 | 11 | 1.00 | 16 | 15 | 0.94 | 14 | 0.88 |
| **average** | | | 0.97 | | 0.67 | | | 0.96 | | 0.68 |
